# *CyAnno*: A semi-automated approach for cell type annotation of mass cytometry datasets

**DOI:** 10.1101/2020.08.28.272559

**Authors:** Abhinav Kaushik, Diane Dunham, Ziyuan He, Monali Manohar, Manisha Desai, Kari C Nadeau, Sandra Andorf

## Abstract

For immune system monitoring in large-scale studies at the single-cell resolution using CyTOF, (semi-)automated computational methods are applied for annotating live cells of mixed cell types. Here, we show that the live cell pool can be highly enriched with undefined heterogeneous cells, i.e. ‘ungated’ cells, and that current (semi-)automated approaches ignore their modeling resulting in misclassified annotations. Therefore, we introduce ‘CyAnno’, a novel semi-automated approach for deconvoluting the unlabeled cytometry dataset based on a machine learning framework utilizing manually gated training data that allows the integrative modeling of ‘gated’ cell types and the ‘ungated’ cells. By applying this framework on several CyTOF datasets, we demonstrated that including the ‘ungated’ cells can lead to a significant increase in the prediction accuracy of the ‘gated’ cell types. CyAnno can be used to identify even a single cell type, including rare cells, with higher efficacy than current state-of-the-art semi-automated approaches.

## Introduction

For many decades, flow cytometry has been used as a conventional technique for both qualitative and quantitative analysis of single cells in a complex cellular system of heterogeneous cell types^1,2^. In recent years, the advancement in the field of single-cell technologies has shifted the paradigm to unravel more complex cell mixtures at a single-cell resolution^3,4^. One such recent technique for single cell profiling at high-throughput level is the Cytometry by Time Of Flight (CyTOF) or Mass cytometry^5^. In CyTOF, heavy metal ion tagged antibodies are bound to single cells, which allow the detection of up to 40 protein cellular markers in millions of cells per sample with much higher sensitivity and reduced “spillover” than traditional flow cytometery^6^. However, the multi-dimensional and complex nature of CyTOF proposes new computational challenge of deconvoluting the heterogeneous mixture of closely related cell types^7,8^.

Conventionally, the deconvolution of a pool of live, single cells into distinct cell types is done by drawing “manual gates” on a hierarchal series of bi-axial plots, wherein the expression profiles of two defined markers are used in a series of sub-setting events to summarize a desired cell population (aka a cell type) with a defined marker expression profile (Figure S1)^9^. The cells that lie inside the defined boundaries of “manual gates” are selected for further sub-selection, whereas, the remaining proportion of cells that remain outside the drawn gates for all cell types are not used in the downstream analysis, i.e. ungated cells (among the non-debris, single, live cells). Here, the choice of gate boundaries is based on expert knowledge by visual inspection and the hierarchical depth of a gating schema depends upon the number of protein markers used in the cytometry panel. For traditional flow cytometry, this process of manual gating is less laborious due to a small number of available parameters^10^. However, as the number of available parameters in CyTOF datasets increases, a deeper interrogation of the cell sub-types can be performed by increasing the hierarchical depth and complexity of the gating schema to reveal previously unknown cell sub-type heterogeneity^11^. This increase in the hierarchical complexity makes manual gating extremely time consuming and laborious, which further increases with sample size^12,13^. As an alternative to laborious manual gating, the most common approach for CyTOF data analysis is the unbiased clustering of the live cells in a high-dimensional space followed by labeling of the cell clusters. The latter is done by analyzing the expression profile for their dominant markers^14^. However, the unsupervised clustering approach suffers from numerous drawbacks that impact downstream interpretation. For instance, random sub-sampling of cells from each sample is routinely carried out to reduce computational complexity and clustering time^15^. Therefore, to overcome some of the limitations associated with unsupervised clustering, semi-automated approaches use “prior” knowledge or “ground truth” about the marker expression in each of the given cell types to annotated every cell of the unlabeled dataset^16^. Currently, only a few semi-automated approaches for cell label predictions are available, viz. Automated Cell-type Discovery and Classification (ACDC)^17^, Semi-supervised Category Identification and Assignment (SCINA)^18^, DeepCyTOF^19^ and Linear Discriminant Analysis (LDA)^10^. Both ACDC and SCINA, uses the list of pre-defined markers for a given cell type to annotate the unsupervised cell cluster(s) that expresses these signature markers. However, these methods depend upon the unsupervised clustering of cells and assume that the expression of target marker as binary (expressed or not expressed), which restricts their ability to classify highly similar cell sub-types, especially non-canonical cell types, that cannot be separated linearly^10,16^. Instead of defined marker lists, DeepCyTOF and LDA use manually gated cell types in form of a marker expression matrix, as training and test set, to build a Machine Learning (ML) model for the cell type prediction. These ML methods can not only automatically learn marker-level features for each cell type, but they also demonstrated high accuracy in classifying non-canonical cell types with small population size. In fact, semi-automated methods like LDA were found to show much higher precision than ACDC and unsupervised methods in correctly predicting cell labels in the pool of gated populations^16^. However, these methods also suffer from elementary limitations, e.g. these methods are built towards the identification of gated cell types only and lack a systematic way for annotating the cells that cannot be classified under any of the desired (gated) cell types, i.e. ungated cells.

In this work, we have shown that the cells that are not assigned to any cell type after manual gating (i.e. ungated cells) are an implicit part of the live cell pool generated by the cytometer and represent a heterogeneous cell population and such cells are difficult to classify as they cannot be explained by the pre-defined set of gating events. Also, the currently published semi-automated methods do not provide a systemic means for incorporating ungated live cells for training, testing and optimizing cell type prediction models. As a result, we have shown that the existing cell annotation methods misclassify these large proportion of ungated cell population as one of the closely related gated cell types.

To address these limitations, we developed a novel semi-automated ML-based computational framework, i.e. CyAnno (*Cy*TOF *Annotator*; Figure 1), that can effectively classify live cells to one of the gated cell types, by learning the marker expression profile of each gated cell population. Our unique approach provides a systematic way for incorporating the marker expression level features of ungated cells for building an optimized ML classification model. Our algorithm demonstrated higher F1 scores, precision and recall rates while differentiating gated populations from each other as well as from ungated populations in CyTOF datasets when the complete dataset with ungated population is taken into consideration during model testing.

**Figure 1.**
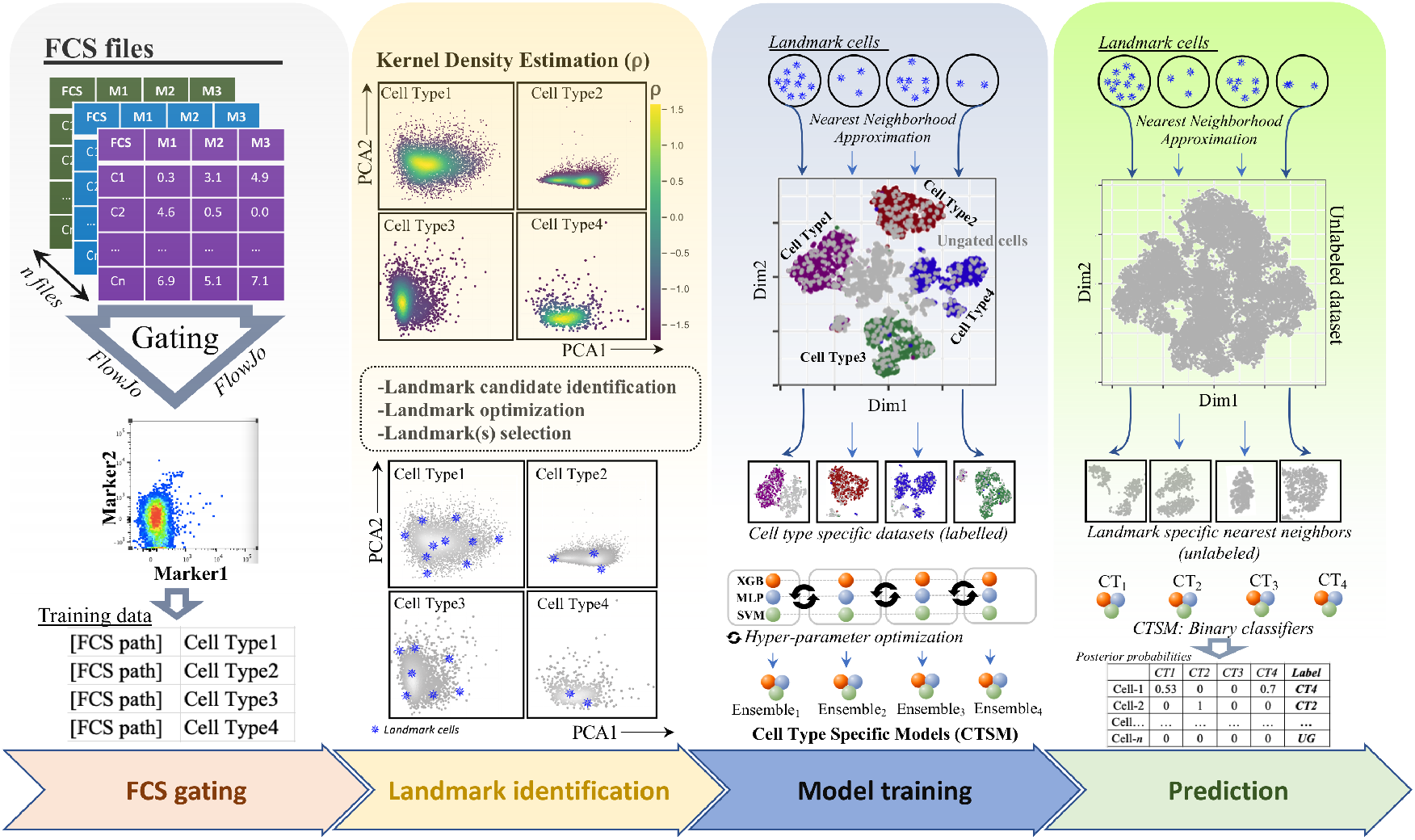
CyAnno Workflow. For details see methods. Briefly, the entire workflow is divided into four steps-*i.) FCS gating*: The first step requires the generation of a training set, which is a collection of CyTOF or flow cytometry FCS/CSV files, one for each cell type manually gated (e.g. by FlowJo) per samples. *ii*.) *LandMark identification*. For each cell type, landmark (LM) cells are identified by computing the kernel densities of each cell in the bi-axial PC plots of its marker expression profile. Candidate LM cells are evaluated using greedy search algorithm to retain only high-confidence representative LM cells (Blue asterisks *). *iii*.) *Model Training*. In the training set with a mixed pool of manually gated cell types, each set of LM cells (one set per cell type) is used to compute their approximated nearest neighbors to create Cell Type Specific Dataset (CTSD). The latter is then used to build a “one-vs-rest” binary ML classification for each cell type, i.e. Cell Type Specific Model (CTSM). The hyper-parameters of the ML models (methods: XGboost, SVM and MLP) are then optimized using random grid search. *iv.) Prediction*. The steps used for CTSD building are applied to the unknown/unlabeled CyTOF/Flow cytometry data. Afterwards, for each cell type, a unique unlabeled CTSD is generated. The final cell labels are predicted by comparing the posterior probabilities of a cell to belong to one of the gated cell types. Cell that do not belong to any of the gated cell type are classified as ‘ungated’ (UG).

## Results

### Characterization of ungated cell population

The analyses of 4 publicly available cytometry datasets (Table 1; see supplementary methods) reveal that a large proportion (~20-60%) of all live cells remain ungated after manual gating (Figure S2). Since every gating event can populate the ungated class of cells, it is expected that together these cells represent a mixture of heterogeneous cell populations. Therefore, when compared with the gated populations in tSNE plots^20^, these ungated cells did not cluster in a defined 2-dimensional space, rather they were visualized as randomly scattered group of cells (Figure 2A). This is further evident by analyzing the silhouette width of each cell type cluster per sample, wherein the cluster of ungated class of cells with very low or negative width indicates their low clustering potential than any of the gated cell types, despite representing a large subset of the live cell population in most of the datasets used (Figure 2B). Here we emphasize that these ungated cells are a not negligible part in a population of live cells and together represent a cell pool with multi-faceted, heterogeneous and undefined marker expression profiles, making their filtration difficult with typical ML approaches.

**Table 1.** Datasets description.

| Table 1. Datasets description. |  |  |  |  |  |  |  |
| --- | --- | --- | --- | --- | --- | --- | --- |
|  | #Samples | #Markers | Mean<br>cell<br>count | #Cell<br>types | Stimulation | #Batches | Reference |
| Levine<br>13 dim | 1 | 13 | 167K | 24 | No | 1 | <sup>25</sup> |
| Levine<br>32 dim | 2 | 32 | 132K | 14 | No | 1 | <sup>25</sup> |
| Samusik | 10 | 39 | 84K | 24 | No | 1 | <sup>26</sup> |
| Multi-center | 16 | 26 | 58K | 4 | No | 2 | <sup>27</sup> |
| POISED | 30 | 39 | 139K | 25 | 2 | 7 | N/A |

For details see supplementary text.

**Figure 2.**
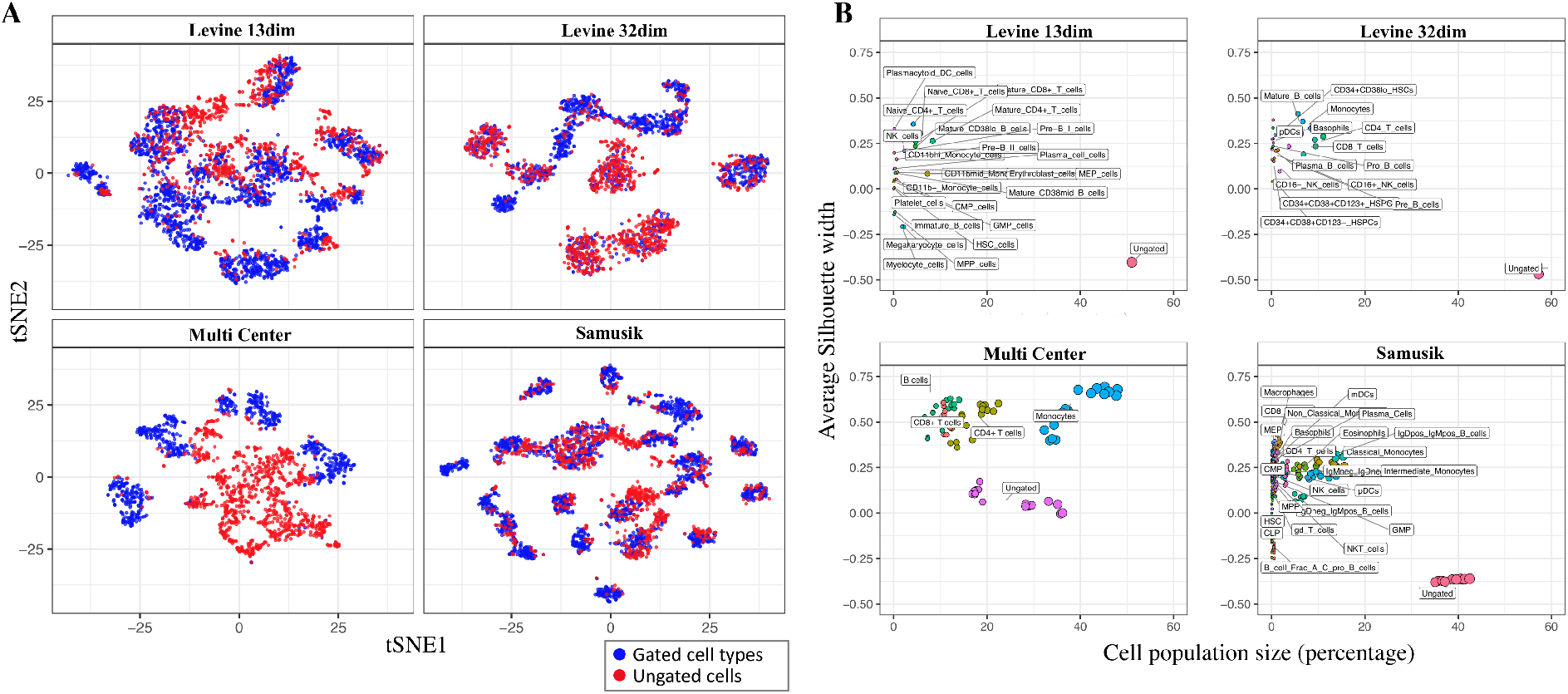
**A.** tSNE visualization to show the unclustered (or random) distribution of ungated cells compared to the gated cell types. **B.** Average silhouette width to show the clustering tendency of different cell types (see materials and methods). In all datasets, ungated cells tend to cluster weaker than any of the gated populations, despite having a large population size in the pool of live cells.

Next, in order to understand the influence of ungated cells on ML model efficacy, we used the published approaches viz. multi-class LDA and deep learning (DeepCyTOF) to build ML models using manually gated cells from the *Samusik* and *Multi-class* CyTOF datasets. These CyTOF datasets have already been used for testing the model efficiency by DeepCyTOF and LDA, however only when the ungated cells were excluded not only from the training but also the test sets^10,19^. Here, we re-evaluated these algorithms on the dataset “with” and “without” ungated cell population in the *Samusik* and *Multi-class* test set. The inclusion of ungated cells in testing sets reflects the real-world scenario in which both the gated population and also the ungated cells are required to be identified. As expected, both LDA and DeepCyTOF predicted class labels with high F1 scores in the absence of ungated cells in the training and test sets, however, the observed efficiency of both models dropped significantly when tested for the prediction ability of these methods on the complete pool of live cells, i.e. gated as well as ungated cell populations (Figure 3 and S3A-B). This is also evident by the posterior probability analysis, wherein we estimated the LDA or DeepCyTOF predicted posterior probabilities of ungated cells to belong to one of the gated populations (Figure 4). The distribution analysis suggests large misclassification of ungated cells to one of related gated cell type with posterior probability > 0.4, a fixed threshold used by these methods below which all cells are considered as unknown/ungated. This leads to the misclassification of ungated cells even with posterior probability > 0.4 to one of the gated cell-types. Alternatively, using a high posterior probability threshold (e.g. 0.7), as suggested by Abdellal et al.^21^, to retain high-confidence prediction and filter out unknown cell populations may not reduce their misclassification, rather it may decrease the overall precision of gated cell type prediction, leading to a higher False Negative (FN) rate. For instance, the predicted posterior probability distribution of many manually gated cell types (e.g. CMP, CLP and plasma cells in the Samusik dataset) is largely centered around a value of less than 0.5 (Figure S4A-B) and when a high posterior probability threshold is used, these cells will be misclassified to the ungated/unknown class of cells. Therefore, increasing the posterior probability threshold might not provide a viable solution to effectively filtering out the ungated cells in the unlabeled live cell dataset while retaining the cells belonging to the gated cell types.

**Figure 3.**
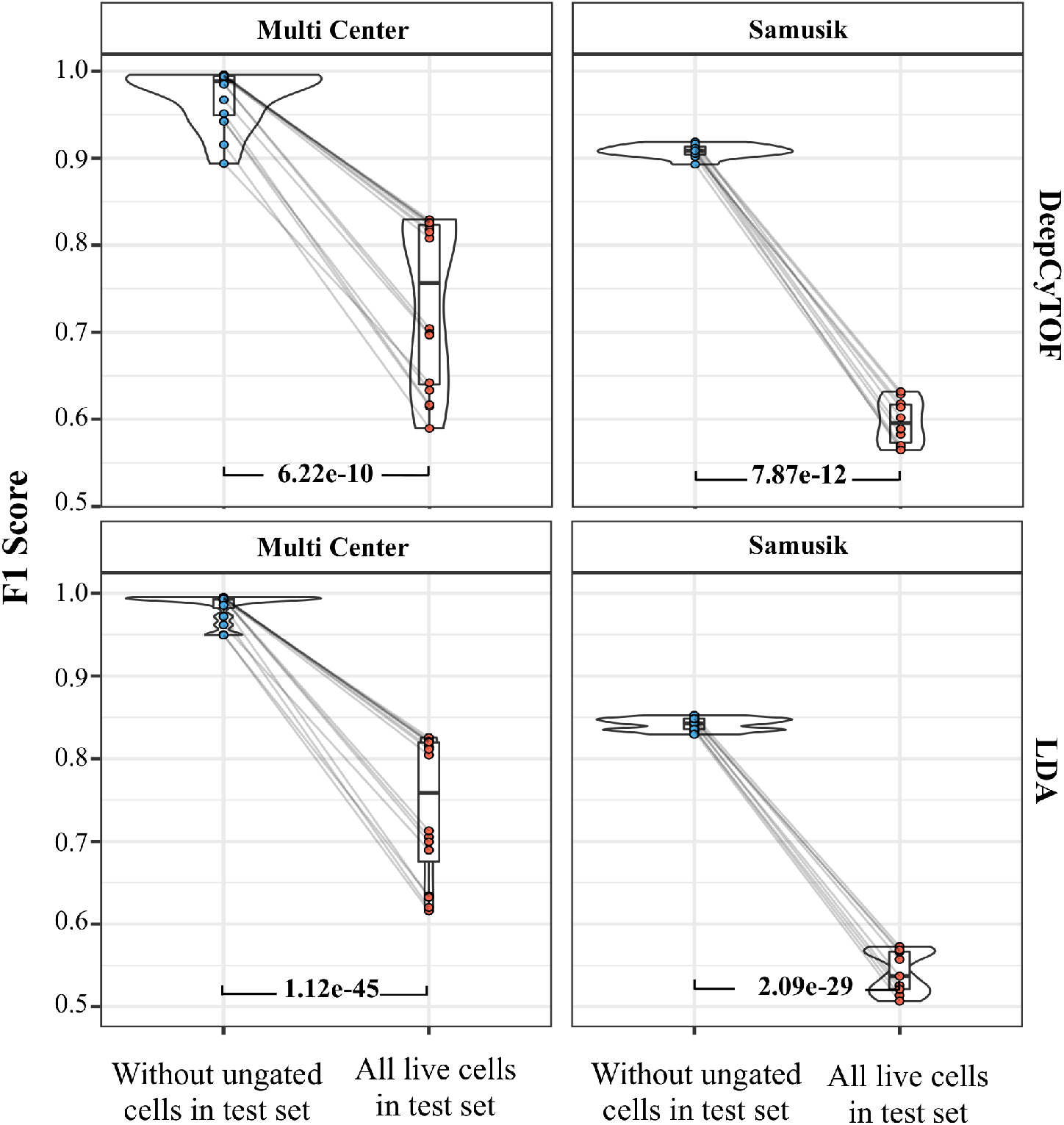
Comparison of per sample F1 score without ungated live cells vs with ungated live cells in the test set after applying DeepCyTOF and LDA to two data sets (Multi Center and Samusik). P-values by paired t-test are shown.

**Figure 4.**
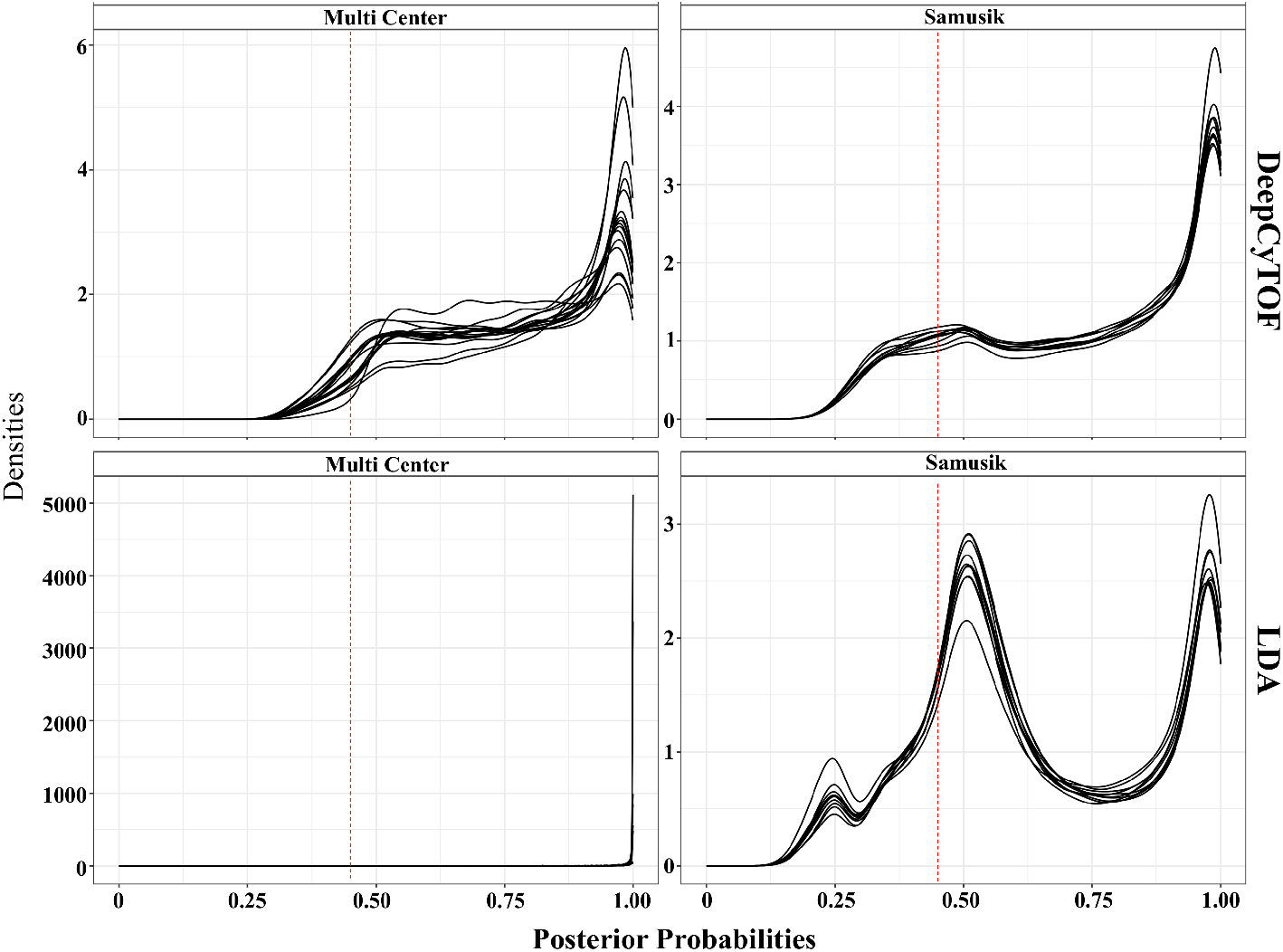
Posterior probability distribution, per sample, of ungated cells predicted by DeepCyTOF (top) and LDA (bottom) to belong to one of the gated populations. Red dotted line shows the probability threshold (0.4) below which cells are classified as ungated by default by DeepCyTOF and LDA.

### Application of CyAnno to the datasets

As one of the initial steps of the CyAnno algorithm (see methods; Figure 1), we identified the LM cells in each of the manually gated cell types in each of the datasets. For each cell type, the proportion of the cells with True Positive (TP) labels that could be retrieved by the LM cells as nearest neighbors is shown in Figure S5, the number of which varies with the proximity of that cell type to the rest of the cell types in a high-dimensional space and the data noise. Wherein, the minimum number of LM cells in each of the cell types is variable and dependent upon the cell type attributes (Figure S6). For canonical cell types with distinct marker profiles, it is expected that only a few LM cells can recall their respective parent cells as nearest neighbors (e.g. in Samusik dataset, only 19 LM cells are required to retrieve 100% pDCs cells from the pool of mixed live cells). The outcome of each LM cell-set neighborhood profiling is the subset of live cells resembling LM cells in terms of marker expression pattern. The resulting neighborhood set is composed of TP labels and FP labels and we called this cell set the “cell type specific dataset”. Figure S7 shows the abundance of different cell types in each of these cell type specific datasets and the ungated cells form, and as expected, the majority of the LM neighborhood cells in most of the cell type specific datasets are FP cells. We used each of the cell-type specific dataset, composed of two class labels- ‘1’ (TP labels) and ‘0’ (FP labels), for building an independent ‘One-vs-rest’ hyper-parameter optimized ML classification model and the subsequent cell type prediction in unlabeled datasets (see methods).

### Comparison with existing methods

Figure S8A-B shows the F1 score achieved by CyAnno with different classifiers for predicting cells of each cell type in the given datasets, wherein XGboost outperforms both MLP and SVM in predicting the correct cell type labels. However, for the evaluation, instead of the single XGboost classifier, we used the consensus of prediction results from all of the classifiers using a majority voting approach to build a cell type specific ensemble classifier. Such Ensemble or hybrid classifiers are less likely to misclassify unseen data and also less probable to over-fit the training set ^22,23^. Therefore, the results of the ensemble classifier in CyAnno were compared to the prediction outcomes of DeepCyTOF and LDA. For both DeepCyTOF and LDA, any prediction with posterior probability less than 0.4 was labelled as unknown/ungated cells, as used in their original publications. The analysis of F1 scores reveals that CyAnno outperforms both DeepCyTOF and LDA when labels of all live cells (i.e. not just the gated population) are used for model evaluation (Figure 5A). For any method, a significant proportion of FP predicted cell type labels belongs to the ungated class of cells, along with other cell types (Figure 5B). Among these predictions, CyAnno performed better than DeepCyTOF and LDA in filtering out the ungated class of cells per sample and simultaneously classifying gated populations with higher efficacy.

**Figure 5.**
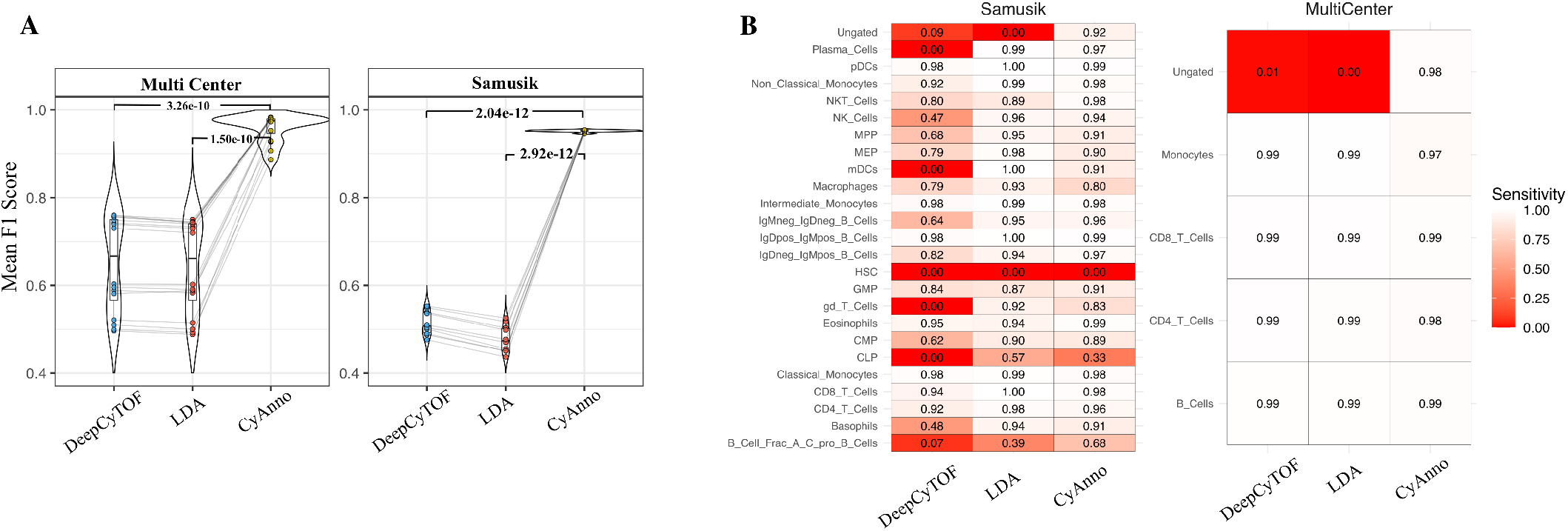
**A.** Mean F1 score comparison (all live cells per sample in test set) with different cell annotation methods tested across two datasets. Mean F1 represents the average of the F1 scores of 5 runs with variable training sets. **B.** The heatmap of prediction sensitivity (=TP/(TP+TN)) associated with each cell type. Here sensitivity highlights the false predictions (red) associated with each cell type. The HSC cell type (in samusik dataset) was found to have very small cell count (< 10) in the training set and, thus, was not used in CyAnno for model training.

Furthermore, to investigate if the inclusion of ungated cells in the training set has played any role in achieving this higher accuracy, we re-executed CyAnno without ungated cells in the training set while keeping them in the test set. We observed that the prediction accuracy dropped significantly in each of the samples if ungated cells were excluded from the training set (Figure 6). The results confirm that the ungated class of cells plays a vital role in guiding the overall performance of models build by CyAnno. Moreover, when the precision score for predicted labels for each cell type was compared with the population size of the respective cell type, CyAnno precision scores were less impacted by the population size than those of DeepCyTOF or LDA (Figure 7). Unlike the other two methods, the CyAnno performed better in predicting each of the cell types with higher precision (minimum precision > 0.75 across all the 5 runs for any cell type) in spite of class label imbalance in the training and test sets. Moreover, the limited variability in the F1 score (across 5 runs; see methods) further suggests the ability of CyAnno in reproducing similar results (Figure S9). Next, for each cell type, we evaluated and compared the composition and proportion of TP and FP cell labels among different cell types predicted by the three different algorithms. As expected, CyAnno outperformed both of the other methods in predicting the highest percentage of TP cells and most of the FP predictions are associated with the ungated class of cells (Figure S10A-B). Whereas, for DeepCyTOF and LDA, the predicted FP cells are composed of numerous gated cell type populations. Ideally, for each cell type, an algorithm should predict low proportion of FP cells with no or minimum number of cells from other gated cell types. For instance, in the Samusik dataset, for NK cell type, the small percentage of CyAnno predicted FP cells are composed of ungated and macrophages only. However, with LDA and DeepCyTOF, FP predicted labels of NK cells are composed of >13 different cell types, which does not include ungated cells in the case of LDA. We observed similar findings for all the other cell types, wherein FP cells predicted by CyAnno are composed mainly of the ungated class of cells. As expected, both LDA and DeepCyTOF fail to identify ungated cells effectively and a very small proportion of ungated live cells are classified correctly. For instance, in the multi-center dataset, LDA fails to identify any of the ungated cell correctly (with predicted posterior probability < 0.4, as used in their original publication) and classified all ungated cells as one of the four gated population (Figure S10B).

**Figure 6.**
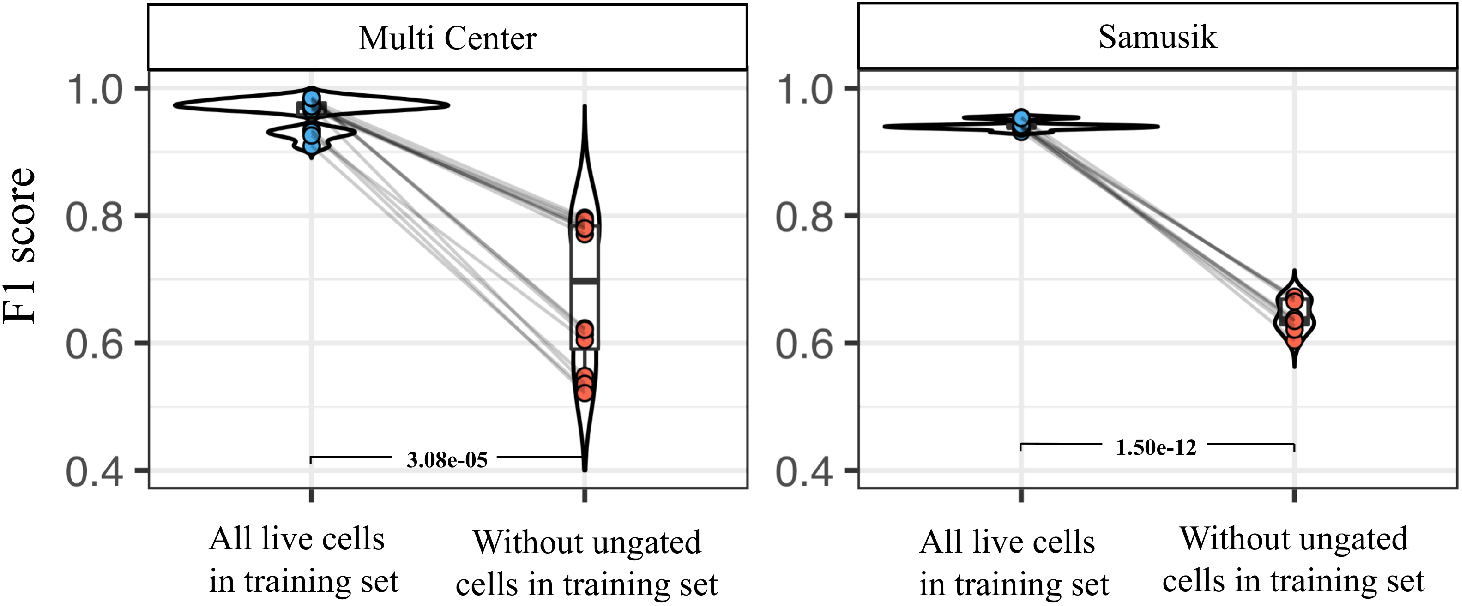
Pairwise sample F1 score comparison of CyAnno when ungated cells are included (all live cells) in model training vs when they were excluded from model training (without ungated cells). P-values were computed with paired t-test, reflecting the statistical significance of difference in outcome when ungated cells are not considered for model training.

**Figure 7.**
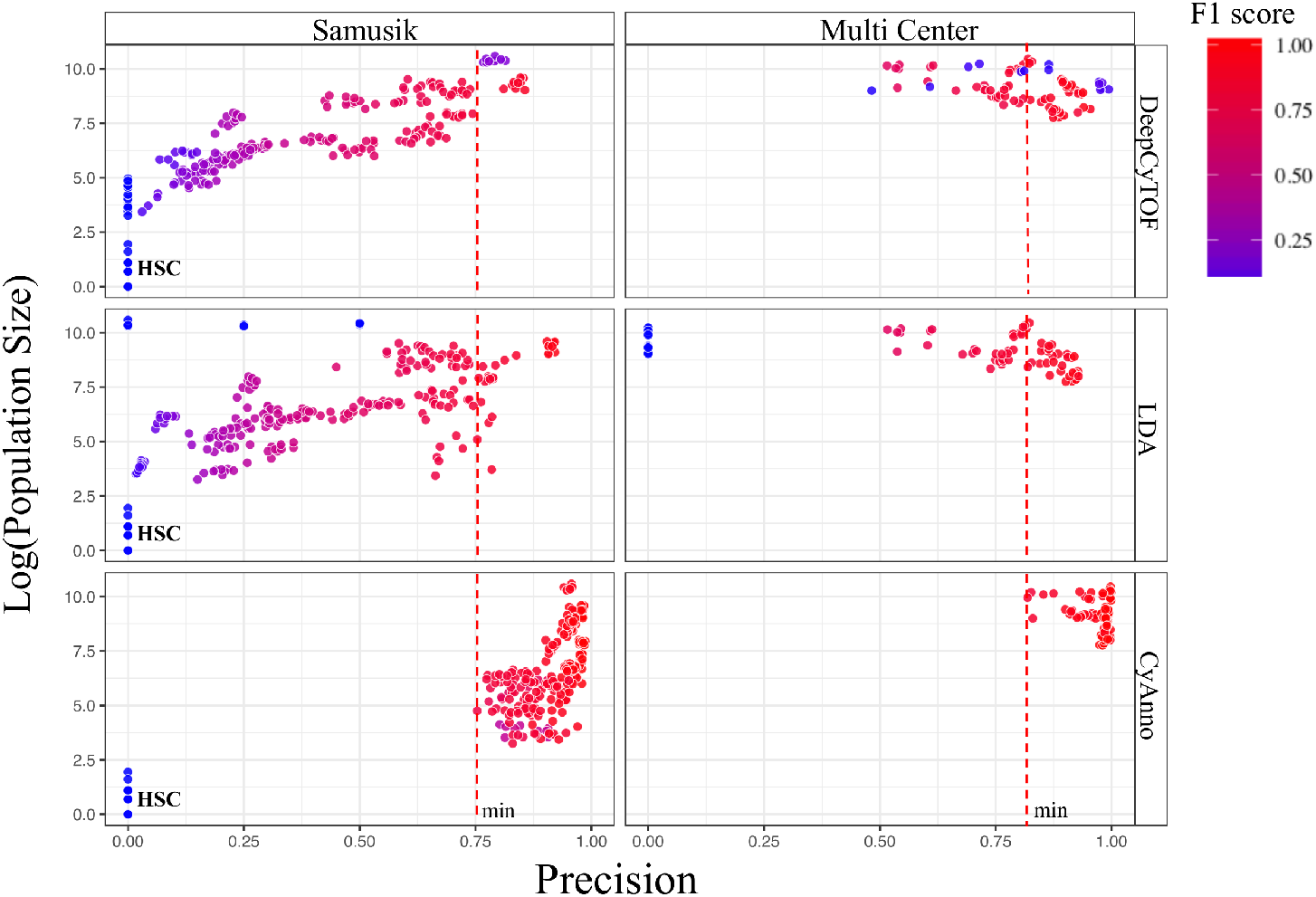
The precision scores (X-axis) are plotted with cell population size (Y-axis; % live cells) of the given cell type in all the samples of the given dataset. Here, the precision score for each cell type was computed using three different methods, each executed 5 times, with varying and independent training dataset in each run. Each circle represents the predicted precision score for a cell type per sample in one run, colored by the F1 score for that cell type during the respective run. The cell type HSC had a very small cell count (< 10 cells) in the overall training set, which did not provide sufficient information for its training, and was thus not included in the training set. The red line marks the minimum precision score observed with CyAnno for the Samusik (0.75; HSC excluded) and Multi Center (0.81) datasets.

### Evaluation with independent benchmark dataset

We also evaluated the performance of CyAnno with an original dataset composed of 15 peanut-stimulated and 15 unstimulated samples, which contains 15-40% of ungated class of live cells (Figure 8A). Out of these 30 samples, we randomly selected 20 independent samples for testing the performance of the three algorithms. From the remaining 10 samples, we randomly selected 3 samples from 3 different batches as training set. For benchmarking with the POISED dataset, we executed each algorithm 5 times with varying training sets (i.e. each time different samples were used) for model building. The mean F1 score, across the 5 runs, per sample in the independent test set was compared and CyAnno outperformed both DeepCyTOF and LDA (Figure 8B). In order to be more certain about the F1 score predicted from CyAnno, we randomly shuffled the cell labels in the test set 1000 times and calculated the p-value of the predicted F1 score (see methods). In all samples, we observed p-value < 0.001 which suggests a very low probability of randomly predicting the observed F1 score.

**Figure 8.**
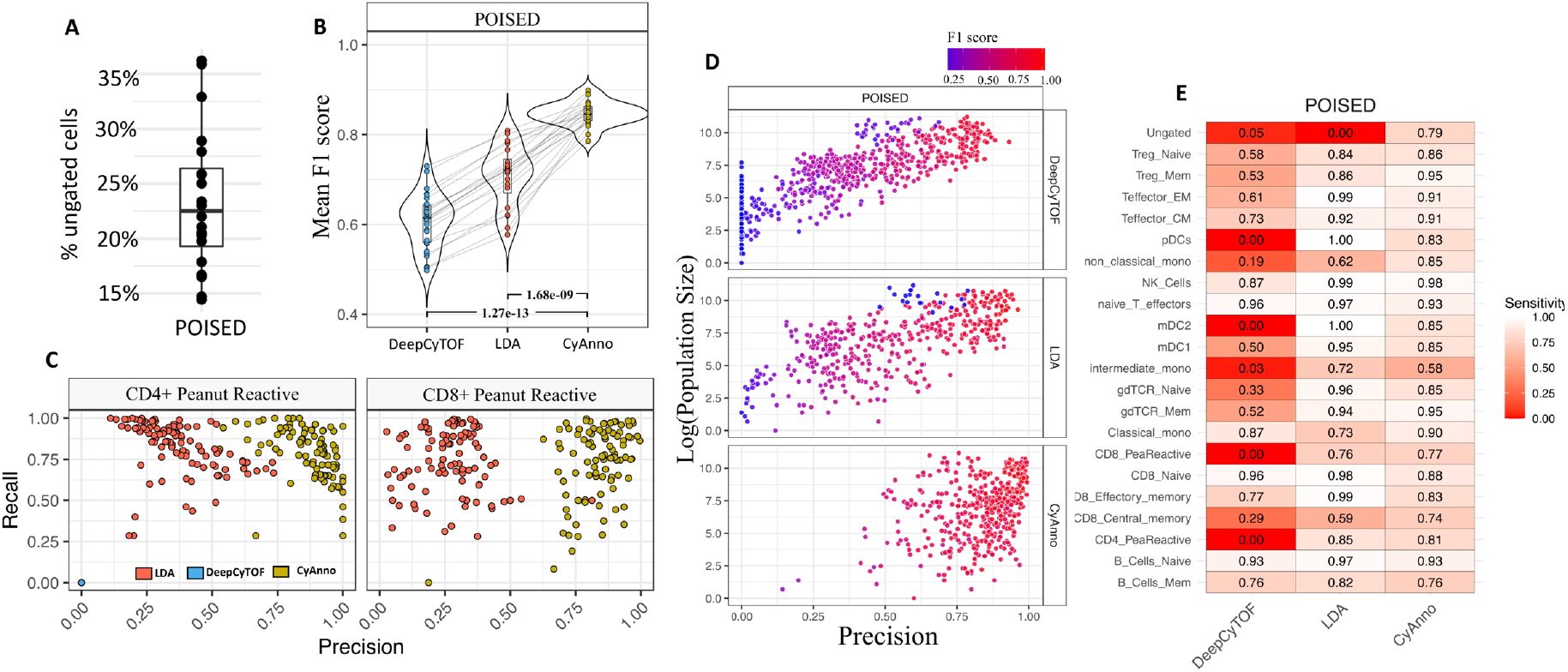
**A**. The percentage of ungated cells per (n=20) files after hand-gating in the POISED dataset. **B**. The F1 score observed with different methods when all live cells are used in POISED training set. **C**. Precision and Recall rate observed for two different peanut reactive cell types with different methods across 5 different runs with varying training sets. Each colored dot represents a predicted score (precision and recall value) observed for the CD4+/CD8+ peanut reactive cell type by three different methods (colored) in the 5 different runs in each test set. CyAnno (yellow colored) predicted labels are found to have better precision than LDA and DeepCyTOF. **D**. The precision scores (X-axis) are plotted with cell population size (Y-axis) of the given cell type in all samples of the given dataset. Here, the precision score for each cell type was computed using three different methods, each executed 5 times, with varying and independent training sets per run. Each circle represents the predicted precision score for a cell type per sample per run, colored by the F1 score for that cell type during the respective run. **E**. The heatmap of prediction sensitivity associated with each cell type. Here sensitivity highlights the false predictions (red) associated with each cell type.

Furthermore, we investigated the overall ability of the different methods to identify CD4+ and CD8+ peanut-reactive cells. Both of these cell types are small in cell population size (proportion < 2% of live cell population) and closely related to many other cell types included in the dataset (Figure S11A). The precision-vs-recall scatter plot analysis revealed the high efficiency of CyAnno in detecting rare cells like these that have a low population size, with high precision and recall rate (Figure 8C). Here, DeepCyTOF failed to predict these cells (recall and precision are 0.0 for all CD4+ and CD8+ peanut reactive cells), whereas LDA predicted these cells with lower precision than CyAnno. Similar to the observation with the other datasets, our analysis reveals that CyAnno’s performance on the POISED dataset is again less impacted by the cell type population size than DeepCyTOF or LDA (Figure 8D). Since, one of the key features of the cell types used in the POISED datasets is their high similarity with each other, we evaluated the composition of FP cell type labels predicted for each cell type (Figure S10C). Clearly, for each cell type, other than TP cell labels, most of the predicted cell labels are composed of ungated class of cells. Whereas, for LDA and DeepCyTOF a large percentage of predicted FP cell labels, for a given cell type, belongs to a diverse set of gated populations. For instance, the FP cells for the CD4+ peanut reactive cell type, when predicted with CyAnno, includes only the ungated class of cells, whereas with LDA and DeepCyTOF ~10 different cell types were falsely predicted as CD4+ peanut reactive cells. Overall, CyAnno has a lower number of false predictions for most cell types, including the ungated class of cells than LDA and DeepCyTOF (Figure 8E).

We also evaluated if CyAnno is capable of predicting only one given cell type from the pool of live cells. For that, we trained the CyAnno algorithm using the CD4+ peanut reactive cell type only (sample size = 3) and predicted these cells in the 20 samples from the independent test set (Figure S12). Similar experiment was also performed for the CD8+ peanut reactive cell type, as both of these cell types have small population size with a mean cell count of 114.17 and 298.78 cells per sample for CD4+ peanut reactive and CD8+ peanut reactive cell type, respectively. The observed results demonstrate the potential of CyAnno in training and predicting even a single cell types with a similar high accuracy. This ability differentiates CyAnno from the rest of the algorithms which essentially require at least two classes to be used for training the ML model(s). Furthermore, we executed CyAnno with varying number of sample size (1 to 10) in the training set, in order to see the effect of sample size on the prediction accuracy with the independent testing set. The analysis shows no drastic change in F1 scores, however a trend for decreased F1 scores can be seen for very small sample sizes (e.g. n=1). For larger sample sizes (e.g. n=10) we observed large F1 scores for all samples in the test set (Figure S13). We next tested if CyAnno predicts biased results for samples that are processed only for a given stimulation, by training the models only from 7 peanut stimulated samples and tested its performance on the 20 independent samples as in previous sections. We compared the F1 scores for the 20 samples by stratifying them with their stimulation status (Figure S14). The results suggest the unstimulated samples were found to have comparatively high F1 score (F1: 0.89-0.99) though only peanut stimulated samples were used in the training set, and the CyAnno results are not biased for the weak stimulations.

## Discussion

CyAnno is a new tool aimed to label the single cells in cytometry datasets by modeling the marker expression profiles through the use of machine learning approaches on manually gated cell populations. We have shown that ‘ungated’ cells are present with a substantially large number within the pool of live cells, after all mutually exclusive series of ‘gating’ events are performed, yet they are largely ignored by the existing cell label prediction approaches. We argued that the primary reason that makes the identification of the ‘ungated’ class of cells difficult is their ambiguous marker expression profile that mimics the expression profile of gated cell types. In fact, in a reduced 2D space (with tSNE), the ‘ungated’ cells are mostly visualized in a cluster with one of the gated cell types (Figure 2). Theoretically, the ‘ungated’ cells can include a ‘unknown’ cell type or cells from any gated cell type just outside the manually drawn gates. The unique feature of the CvAnno algorithm is that it explicitly learns the boundaries between ‘ungated’ cells and the gated cell types in a high-dimensional space and then predicts the cell labels in the unlabeled samples. By accounting for the ‘ungated’ class of cells in the datasets, we have shown that CyAnno can predict cell labels with much higher accuracy than existing approaches. In particular, our analysis shows that ignoring the ‘ungated’ cells during model training can lead to less specific and inaccurate identification of the gated cell types. This information is not used by existing (semi-)automated computational approaches while annotating unlabeled CyTOF datasets, as they ignore ‘ungated’ cell annotation during model training and its performance evaluation. For the other semi-automated methods, using ungated cells for modelling the cell type is a technical challenge, as these cells represent a heterogeneous population of cells from a mixed cell type and together cannot be used as a single class of cells. Therefore, the simplest solution to predict the unknown cells adopted by currently available semi-automated approaches, as suggested by Abdellal et al.^21^, is to increase the posterior probability threshold (e.g. > 0.7) to recall only the ‘gated’ cell type with higher confidence. However, we found that that this would significantly increase the number of ‘False Negative’ predictions as a large proportion of gated cells may have predicted posterior probabilities of < 0.7. CyAnno overcomes this problem by building a cell type specific ‘one-vs-rest’ binary model, in which the predicted posterior probability for a TP cell to be assigned to its cell type by the respective model should be > 0.50 and it is independent of the other cell types as well as ‘ungated’ cells.

In addition, CyAnno also performed exceptionally good in the identification of CD4+ and CD8+ peanut-reactive cell types which are very small in terms of population size but highly relevant in peanut allergy research. Such small populations are generally not captured using unsupervised approaches and also the other two semi-automated algorithms used in this study failed to reach similar level of high precision as compared to CyAnno. However, the prediction accuracy of the algorithm may also depend upon the choice of training set, and we recommend the inclusion of samples processed under different batches or stimulation for training the models, to keep the overall training unbiased for any given batch or stimulation. However, the stimulation used here was weak and the performance of the algorithm has not been tested on samples with strong stimulation that leads to aberrant marker expression profile.

One of the other factors that can affect the prediction accuracy of any cell type is the ambiguity in its marker expression profile as compared to the other cell types. For instance, pDCs, in both POISED and Samusik datasets, express a unique expression profile and therefore visualized as distinct cluster in tSNE plots (Figure S11A and S11C). Such cell type, irrespective of the population size, not only required small number of LM cells to be retrieved from the pool of live cells with mixed cell types (Figure S6) but their predicted F1 score was also found to be high. However, other cell types can closely resemble each other in a high-dimensional space (e.g. naïve B cells vs memory B cells in the POISED dataset). Such cells are often difficult to classify and require a higher number of LM cells. Therefore, the degree of similarity of cell types used for classification can play a significant role in the prediction accuracy of final models. Since, the cell types used in both Samusik and Multi-Class datasets are well defined in their marker expression profile and are more distinct (Figure S11B-C) than the cell types used in POISED (Figure S11A), the latter stands as a valuable resource for benchmarking the efficacy of prediction models on CyTOF datasets.

One unique application of CyAnno is its ability to train and predict even a single cell type, irrespective of its population size (Figure S12). This feature differentiates the application of CyAnno from the rest of the existing algorithms. The other algorithms typically require multiples classes (i.e. cell types) for training the model in which the predicted posterior probability of a cell to belong to a given cell type is computed with respect to the other cell types. In contrast to that, in CyAnno, each cell type is trained as a separate one-vs-rest model, therefore, the overall predicted posterior probabilities of the cells of a given cell type are independent of the rest of the cell types. This essentially allows CyAnno to train for and predict even a single cell type from the pool of unlabeled live cells with a mixed cell type population. This unique feature of CyAnno can be useful for a broad range of studies; e.g. when only one (or few) cell types are of clinical interest, especially those rare cell types that cannot be clustered with unsupervised clustering. CyAnno can be used in studies in which a specific cell type (e.g. as shown for CD45+ Lin^lo^ cells in Hamers et al.^24^) is manually gated and then exported for downstream analysis. This can save efforts and time during manual gating and can assist focused research without compromising on overall accuracy.

One of the known limitations of CyAnno is its higher computational cost than DeepCyTOF and LDA (Figure S15, see supplementary text), wherein hyper-parameter optimization (for all of the ML algorithms) consumes the majority of the computational time. Since, the hyper-parameter optimization is the most time consuming yet unavoidable part of CyAnno, the future plan is to incorporate Bayesian hyper-parameter optimization that can evaluate much a larger combination of hyper-parameters with lower computational cost.

## Methods

### Datasets

In order to perform comparative analysis with existing approaches we used publicly available and one previously unreported CyTOF datasets for algorithm development and evaluation, viz. Levine 13 dim^25^, Levine 32 dim^25^, Samusik^26^, Multi-center datasets^27^ and POISED (Table 1). The details of different datasets used along with experimental details and other information for generating the POISED CyTOF dataset are available in supplementary text.

### CyAnno Algorithm overview

Here we propose a unique semi-automated approach, viz CyAnno, for classifying cells to one of the gated cell types or the ungated cell population (Figure 1 and Figure S1). The input was a training set, which can be a collection of CyTOF or flow cytometry FCS/CSV files, one for each cell type *m* manually gated per samples and used for training, with marker expression profiles 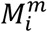 of live cells along with ungated class of cells *U* from each sample *i* (non-debris; non-doublets). The method leveraged prior knowledge about the marker-level features expressed by the ungated cells U as well as the cells of each of the manually gated cell types *m* in each of the sample *i* of the training set. The result of the algorithm was an independent “one-vs-rest” classification model, one for each cell type.

### CyAnno workflow

Each CyTOF sample S*_i_* was pre-gated to include millions of non-debris, non-doublets single cells *n_j_* in which each cell *j* is an independent measurement event with expression values of *p* markers, which can be denoted as *n-by-p* matrix. The aim was to stratify the cells that had similar expression of the set *p* markers, i.e. cells that belong to the same cell type *m*, where 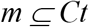. In order to label each cell to a given cell type, we performed manual gating (e.g. using FlowJo) and the expression matrix 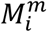 for each of the gated cell population were exported from each CyTOF sample *i*. Each sample S*_i_* also contained cells that were not assigned to any of the cell types during manual gating. We call these cells ungated or unknown, and they represented a population of diverse or mixed cell types. Thus, if *U* denote the unknown/ungated cell population then the total live cell population of each sample *i* can be represented as *S_i_* ∈ {*Ct*, *U*}. Overall, the CyAnno was implemented as a 3-step serial framework:

1. *Identification of Ungated cell*: For each training cytometry sample *S_i_*, manual gating was performed and each of the gated cell types is exported into a separate FCS file, i.e. one FCS file per cell type per training sample. Thereafter, we identified the cells that were not part of any of the gates, i.e. ungated cell population, by comparing the cells of the gated cell populations with the pool of live cells (Figure S1).
2. *Data Transformation*: The marker expression profile of gated cell types and ungated cells were then transformed using arcsinh transformation (co-factor set to default value of 5; user-defined) to facilitate downstream clustering and ML model building. The transformation provided necessary scaling to build efficient ML models and minimizes the inherent noise associated with marker expression values.
3. *Building the ML models from training set*: The following details the five sub-steps in CyAnno workflow that are performed for each gated cell type from a set of *Ct* cell types:
  a. *Landmark cell identification*: Within each of the manually gated and mutually exclusive cell type expression matrixes 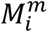, we identified a small set of characteristic cells called LandMark cells (LM cells). The essential feature of the LM cells was their ability to effectively identify the cells of its respective cell type as nearest neighbors in a population of mixed cell types and ungated cells. The rationale behind this was that the LM cells in a high-dimensional space should closely resemble the cells of its corresponding cell type. Therefore, the nearest neighbors of the LM cells in a pool of mixed cell population were expected to include most (or all) of the cells from their respective cell type (i.e. TP cells) along with cells of closely related cell types, i.e. FP cells. We applied a stringent protocol to identify high-quality LM cells from each of the gated cell types that can retain maximum number of TP cells. The following approach was applied to shortlist LM cells for each cell type:
    i. *Decomposition into Principal Components (PCs)*: The manually gated multi-dimensional expression matrix 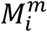 is decomposed into its first two Principal Components (PCs), i.e. PC1 and PC2.
    ii. *Kernel Density Estimation*: In order to identify LM cells, kernel densities within the first two PCs were estimated. The kernel density values smooth out the contribution of each data point over its nearest neighborhood in PCs biaxial plots, where the estimated density at each point can be represented as:

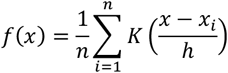 Where estimate of *f*(*x*) is the convolution of Gaussian Kernel *K* (mean value of 0) with the 2D histogram of PCs and *h* is the bandwidth estimated from data points.
    iii. *Landmark candidates shortlisting*: We then stratified the estimated kernel densities into its 5 quantile ranges and randomly select 10 cells in each of the quantiles as LM candidates. This allowed us to shortlist a restricted set of LM candidates as potential representatives of the cell type. In addition, cells that form the edges in the PCs biaxial plots were estimated by computing the alpha shapes (method: concave hall; alpha = 0.80) of a set of points. The latter cells represented highly diverse cells of the cell type to be included in the set of LM candidates.
    iv. *Landmark cell selection via nearest neighborhood approximation*: Using the Nearest Neighborhood Approximation (NNA) via Facebook’s similarity search and clustering algorithm, viz faiss (https://github.com/facebookresearch/faiss), we identified the nearest neighbors of each LM candidate within all live cells. This method performed the nearest neighbor search using a brute-force index with L2 distances. Here, the number of nearest neighbors to be searched for each LM candidate was equivalent to the proportion of the given cell type population in the total live cells. Ideally, the LM candidate with highest kernel density will enumerate the largest number of cells belonging to their respective cell type, i.e. TP cells, however, when a cell population contains highly similar cell types, each LM candidate can produce lower than the expected proportion of TP cells. Therefore, the algorithm shortlisted and prioritized the LM candidates using the greedy search algorithm, which optimized for smallest number of LM cells that could retrieve the highest number of TP cells. Starting with LM candidate cells with the highest kernel density, we selected the total number of LM candidates required to capture >99.99% of TP cells as nearest neighbors form the LM cells, such that no two LM cells shared more than 20% of the same TP cells. If two LM candidates shared more than 20% of nearest neighbors, the LM cell with a larger number of TP cells is selected. However, if the total number of cells in a given cell type was ≤ 100 (default value, user defined in CyAnno wrapper) and predicted TP nearest neighbor cells were < 99.99% then all cells for that given cell type were considered as LM cells.
  b. *Generating Cell Type Specific Training Dataset*: For all selected LM cells, the nearest neighbors (by NNA) within all live cells of the training set was composed of a sub-set of cells with true labels for the given cell type (i.e. TP cells) along with the nearest neighbor cells that did not belong to the cell type (i.e. FP cells). We called each of this cell subset of nearest neighbors a “Cell Type Specific Training Dataset”.
  c. *Building a Binary Cell Type Specific Model* (*CTSM*): Each “Cell Type Specific Training Dataset” was composed of two kinds of labels- “0” for FP nearest neighbors and “1” for TP nearest neighbors. This training dataset was used to build the ML-based binary classifier(s). CyAnno used three different ML algorithms for building prediction models, i.e. extreme gradient boosting (XGboost)^28^, Multi-Layer Perceptron (MLP)^29^ and Support Vector Machine (SVM)^30^. For each of the ML algorithms, the hyper-parameters were optimized using random grid search with cross validation analysis, wherein the large search space was defined for different parameter options for each ML algorithm.
  d. *ML parameter tuning and error estimation* For XGboost, decision tree algorithm with base learner “*gbtree*” was used with learning task objective set to “*binary:logistic*” for binary classification per cell type. For random grid search space, the set of gamma ∈ {0.5, 1.0, 1.5, 2.0, 5.0, 10}, validation subsampling ∈{0.3, 0.2} and the learning rate ∈{0.01, 0.1, 0.3} were used with maximum tree depth of 6. For feature selection, “*thrifty*” selector with setting of top 5 features per group was used, with L1 regularization weight (*<u>reg_alpha</u>*) set to 0.005. Models were evaluated using validation dataset with two different evaluation matrices-logloss matrix and binary classification error rate (#FP / #(FP+TP)) in which the cell label prediction score larger than 0.6 were used as positive instances. Usually, the prediction score of >0.5 for the binary classification should be sufficient, however, the score of 0.6 ensures for higher precision in outcome. Early stopping parameter was also used, in which model training terminates if validation score did not improve in 10 consecutive iterations. For MLP, Stochastic gradient descent (*sgd*) solver was used for weight optimization with adaptive learning rate. For hyper-parameter optimization, the sets of L2 regularization term ∈ {1, 0.1, 0.01} and initial learning rate ∈{0.01, 0.1, 1.0} were used with validation subsampling ∈ {0.3, 0.2}. Early stopping parameter was set to 10 which terminated model training if validation score did not improve by value of 0.001 in 10 consecutive iterations. For SVM classifier, Radial Basis Function (*rbf*) kernel trick was used for building non-linear classification boundaries. Here, hyper-parameters set of C∈{0.1, 0.01, 0.001} and γ ∈{1, 0.1, 0.01} were used for balancing SVM classification error, regularization and bandwidth estimation. In all of the cases, a total of 10 random hyper-parameters combinations were used for model training and evaluation. With the CyAnno wrapper, these algorithms can be called independently or they all can be used together for predicting cell labels using an ensemble model, which combined the results of multiple base estimators and provide consensus results by majority voting approach. For the subsequent analysis in this work, ensemble of ML models was used for predicting cell labels.
4. *Cell type Classification with a CTSM*: To annotate a new data set (e.g. validation dataset or new dataset) of unlabeled live cells the following steps were performed per cell type (*m*): First, NNA was performed using the previously predicted LM cells for the cell type. The resulting “cell type specific dataset” was then applied to its corresponding CTSM built previously (from section *c*). For each cell type *m*, the analysis of unlabeled dataset using the CTSM results in a posterior probability *p* of each of its cell *j* belonging to the given cell type *m*, such that

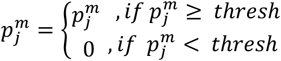 Where 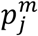 was the predicted posterior probability of a cell *j* belonging to a cell type, *m*, in a binary classifier. This predicted posterior probability becomes 0 if it was less than user defines threshold, i.e. *thresh* (default 0.5; user-defined). The final posterior probability matrix 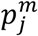 contained the probability score of each cell *j* of belonging to each of the cell type *m* of the set *Ct*. From the probability matrix, the final cell type label *l* for each cell *j* was defined by:

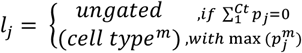 Where, max 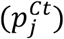 is the maximum posterior probability observed for a given cell *j* across all the cell types. The final cell type label of a cell *j* was assigned as a cell type *m* with maximum posterior probability.

### Comparison with existing methods

To evaluate the efficacy of CyAnno, we applied the proposed pipeline to the Samusik, Multi-Center and POISED CyTOF datasets. The three methods (CyAnno, DeepCyTOF, LDA) were tested across five different runs, in which, during each run, 20% of the randomly selected samples were used for training the model (i.e. training set), whereas the rest of the samples were evaluated for model testing (i.e. test set). The resulting F1 scores across the runs for each sample as well as for each of the given cell type were then compared for evaluating the prediction ability of the three different methods. For CyAnno, we used ungated cells in the training set for cell type classification, unlike the LDA and DeepCyTOF methods which did not allow the inclusion of ungated cells for model training.

### Performance matrices

a. The CyAnno performance was compared and evaluated in terms of F1 score which was defined as:

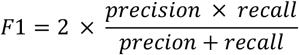 Where, *precision* = *TP/TP* + *FP* and *recall* = *TP/TP* + *FN*.
b. To further confirm the accuracy of built models in predicting correct cell labels, we also performed a permutation test of predicted labels to calculate the significance of predictions. Wherein, the classification accuracy of CyAnno was compared after randomly permuting the cell label *1000* times and examining the change in sample F1 score. The statistical significance was measured by *p* value which reflected the probability of obtaining a high F1 by chance, and calculated as:

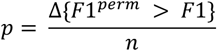 Where, Δ is the cardinality of set when the F1 scores of permuted samples were larger than or equal to the F1 score obtained with original sample cell labels. Low *p* value (e.g. p < 0.05) suggest high accuracy in correctly predicting cell labels.

### CyAnno implementation

All codes were compiled with Python programming language (version 3.0 or above) on Linux platform with sckit-learn, NumPy and Pandas libraries. A Lenovo P400 workstation running on 2 X Intel-Xeon processor with 20 cores and 64 GB of RAM was used. Although CyAnno has been tested only on Linux based OS, including MacOS, however, for the end-users, the CyAnno algorithm is wrapped as an easy-to-use python script that can be executed via command line interface on any OS supporting python.

## Data availability

The publicly available datasets can be obtained from HDCytoData R package^31^. Multi-Center dataset can be obtained as publicly available datasets from GitHub^10^ (https://github.com/tabdelaal/CyTOF-Linear-Classifier). Raw unlabeled POISED dataset files in FCS 3.0 format and normalized labelled CSV format will be available after publication.

## Code availability

The CyAnno is freely available as an easy-to-use python script with a user-manual and sample dataset at https://github.com/abbioinfo/CyAnno.

## Acknowledgement

This work was supported by the National Institute of Allergy and Infectious Diseases (NIAID) under grants 5U19 AI104209-07, R01AI140134-01 and 5U01 AI140498-03, the Sean N Parker Center for Allergy and Asthma Research at Stanford University. Also, this work was partially supported by the National Institute of Health Grant P30-CA124435, Grant P30-DK116074, and Grant UL1-TR003142.

