## supplementary for "*CyAnno*: A semi-automated approach for cell type annotation of mass cytometry datasets"

### Supplementary information

**Supplementary Table 1**. Panel of metal-conjugated antibodies used for mass cytometry analysis of PBMCs for POISED dataset.

| Marker | antibody | Marker type |
|---|---|---|
| CD19 | Nd142Di | Lineage |
| CD49b | Nd143Di | Lineage |
| CD4 | Nd145Di | Lineage |
| CD8 | Nd146Di | Lineage |
| CD20 | Sm147Di | Lineage |
| CD38 | Nd148Di | Lineage |
| CCR4 | Sm149Di | Lineage |
| LAG3 | Nd150Di | Lineage |
| CD123 | Eu151Di | Lineage |
| CD45RA | Eu153Di | Lineage |
| CD3 | Sm154Di | Lineage |
| HLA-DR | Gd157Di | Lineage |
| CD33 | Gd158Di | Lineage |
| CD11c | Tb159Di | Lineage |
| CD14 | Gd160Di | Lineage |
| CD69 | Dy162Di | Lineage (To identify pea-specific cells) |
| CXCR3 | Dy163Di | Lineage |
| CD127 | Ho165Di | Lineage |
| CD27 | Er167Di | Lineage |
| CD40L | Er168Di | Lineage (To identify pea-specific cells) |
| CCR7 | Tm169Di | Lineage |
| CD25 | Yb173Di | Lineage |
| CD56 | Yb174Di | Lineage |
| TCRgd | Lu175Di | Lineage |
| CD16 | Bi209Di | Lineage |
| CD86 | In113Di | Functional (Not used in analysis) |
| OX40 | Pr141Di | Functional (Not used in analysis) |
| CD28 | Gd155Di | Functional (Not used in analysis) |
| GPR15 | Gd156Di | Functional (Not used in analysis) |
| PD1 | Er170Di | Functional (Not used in analysis) |
| beta 7 integrin | Yb172Di | Functional (Not used in analysis) |
| CLA | Yb176Di | Functional (Not used in analysis) |
| CD57 | Y89Di | Functional (Not used in analysis) |
| IL-4 | Nd144Di | Functional (Not used in analysis) |
| IL9 | Sm152Di | Functional (Not used in analysis) |
| IFNg | Dy161Di | Functional (Not used in analysis) |
| IL-17 | Dy164Di | Functional (Not used in analysis) |
| LAP | Er166Di | Functional (Not used in analysis) |

| | | |
|---|---|---|
| IL-10 | Yb171Di | Functional (Not used in analysis) |
| * Only Lineage markers were subjected to CyAnno for model training and cell type identification. | | |

**Dataset description**

Overall 5 datasets were used in this study for method development and result evaluation, viz. Levine 13 dim[1], Levine 32 dim[1], Samusik[2], Multi-center datasets[3] and POISED (Table 1).

The Levine 13 dim is a CyTOF dataset in which samples were taken from human bone marrow cells of a single healthy donor, with a panel composed of 13 protein markers and 24 manually gated cell types. Similarly, Levine 32 dim is a CyTOF dataset composed of 32 protein markers with 14 manually gated cell types from human bone marrow cells from two healthy donors[1]. The "samusik" (or PANORAMA) CyTOF dataset was obtained from Samusik *et al.*[2] and contains samples from bone marrow of 10 different mice in which the expression profile is measured using a panel of 39 markers. The "Multi-center" dataset contains 16 samples and a panel of 26 markers in which samples were run across two different centers (or batches)[3]. While the samusik dataset contains data of 24 different cell populations, the multi-center dataset contains only 4 cell types. In the samusik dataset, the cell type HSC had a very small cell count (< 10 cells; mean ~3 cells per sample) in the overall dataset and were therefore excluded from the training set, however, the HSC cells remained as the part of the test set. The publicly available CyTOF datasets used in this study were obtained from the respective publications or HDCytoData R package[4]. Despite being commonly used in many studies, we found that these and other public datasets with manually gated cell type labels have one or more of the following limitations for testing any cell type prediction algorithm. The shortcomings include small sample size, small number of batches (≤ 2), mostly distinct with only few closely related cell types, and no varying biological treatments. In fact, the limited sample size did not allow us to perform exhaustive model evaluation on large independent datasets belonging to untrained batches or stimulations. Therefore, we relied on an additional dataset (called POISED[5]) of 30 CyTOF samples from human PBMCs, from peanut-allergic individuals, for testing our algorithm. These samples were run across 7 batches with a panel of 21 lineage markers and 18 functional markers (Table S1) and under two different stimulations, i.e. unstimulated and peanut stimulated. 21 cell types were manually gated (including closely related cell types). The non-canonical peanut reactive T cells used in this study were CD69+ CD40L+ CD4+ T cells and CD69+ CD8+ T cells [6].

**Computational Time for different methods**

Since four different CTSMs can be built for each cell type, the total computational cost for running CyAnno is directly proportional to the number of cells, cell types and ML algorithm used. Figure S15 depicts the total time taken to build an optimized CTSM based on XGboost, MLP and SVM for each cell type in the POISED dataset. Here most of the time is used in hyper-parameter optimization which is an essential part of any ML based pipeline. For faster results, we found that XGboost outperforms any of the other ML algorithms used. Although, for highest accuracy we recommend using ensemble classifier as it uses prediction results from multiple ML algorithms and thus less likely to produce over-fitted results

### Supplementary Figures

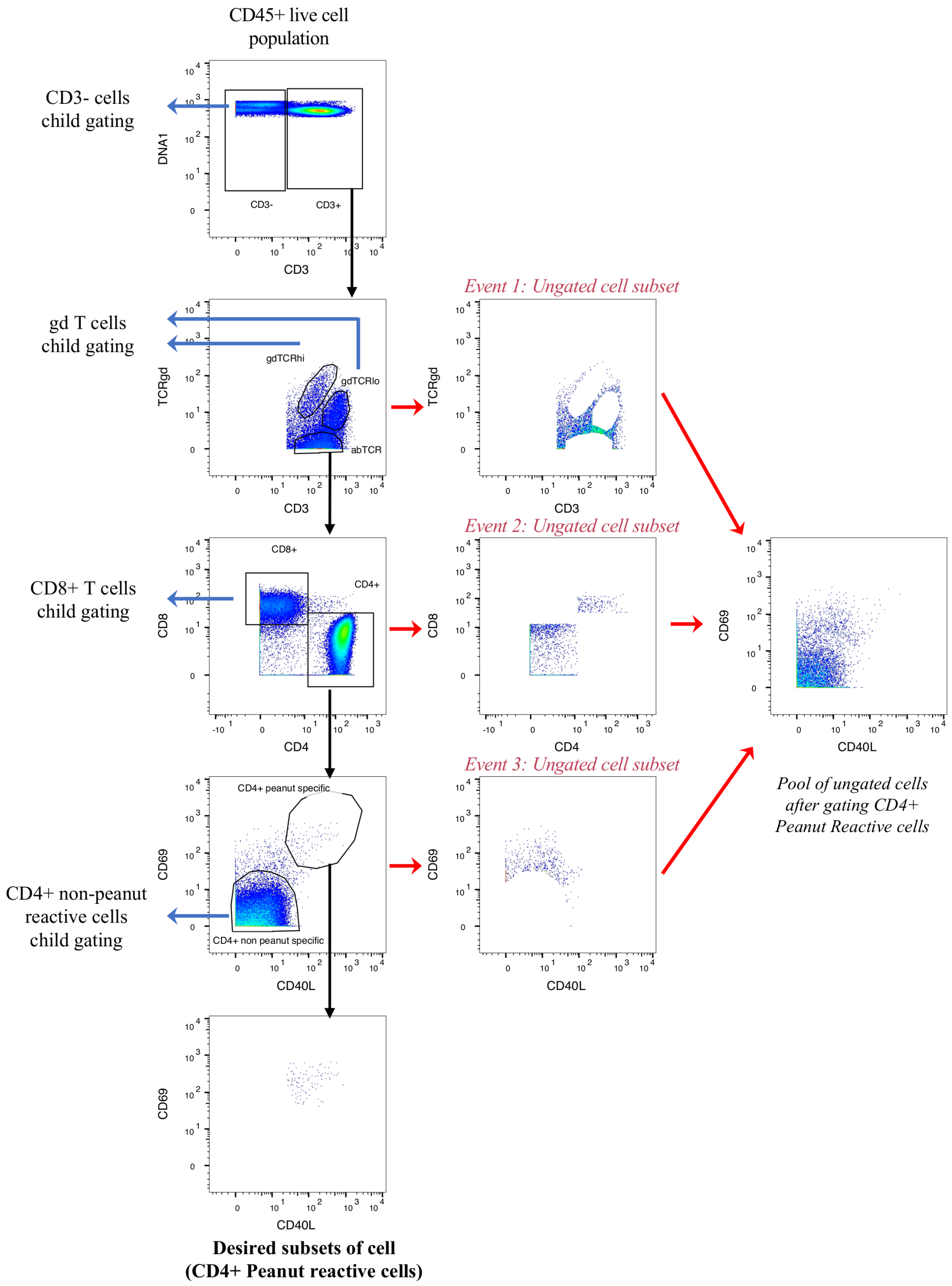


**Figure S1**. Manual gating of CD4+ Peanut reactive cell type. The hierarchical schema illustrates the process involved in identification and gating of CD4+ Peanut reactive cells. Each gating event discards a population of cells as undefined, i.e. ungated. It is to be noted that each gating event can further lead to a hierarchical series of downstream child-gating for identification of other cell types (highlighted on the left). Therefore, once all the mutually exclusive cell types are identified, these ungated cells together form a major population composed of heterogeneous pool of live cells, that are expected to be detected as ungated class of cells by (semi-)automated approaches

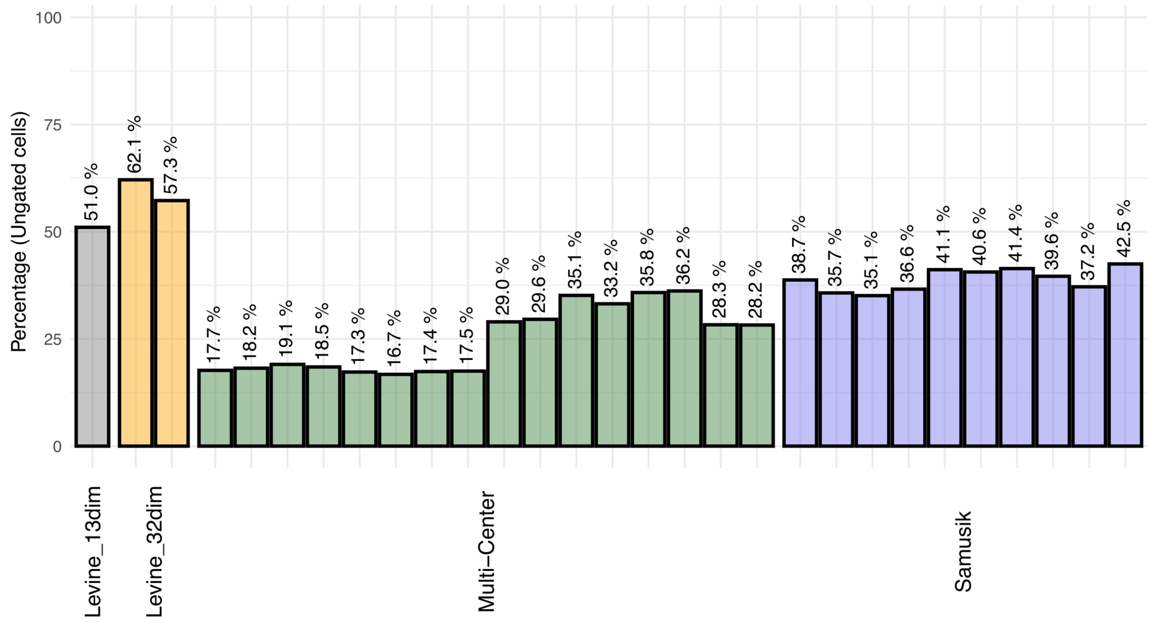


**Figure S2**. Percentage of ungated cells within live cells, after manual gating, in each sample of the 4 public datasets.

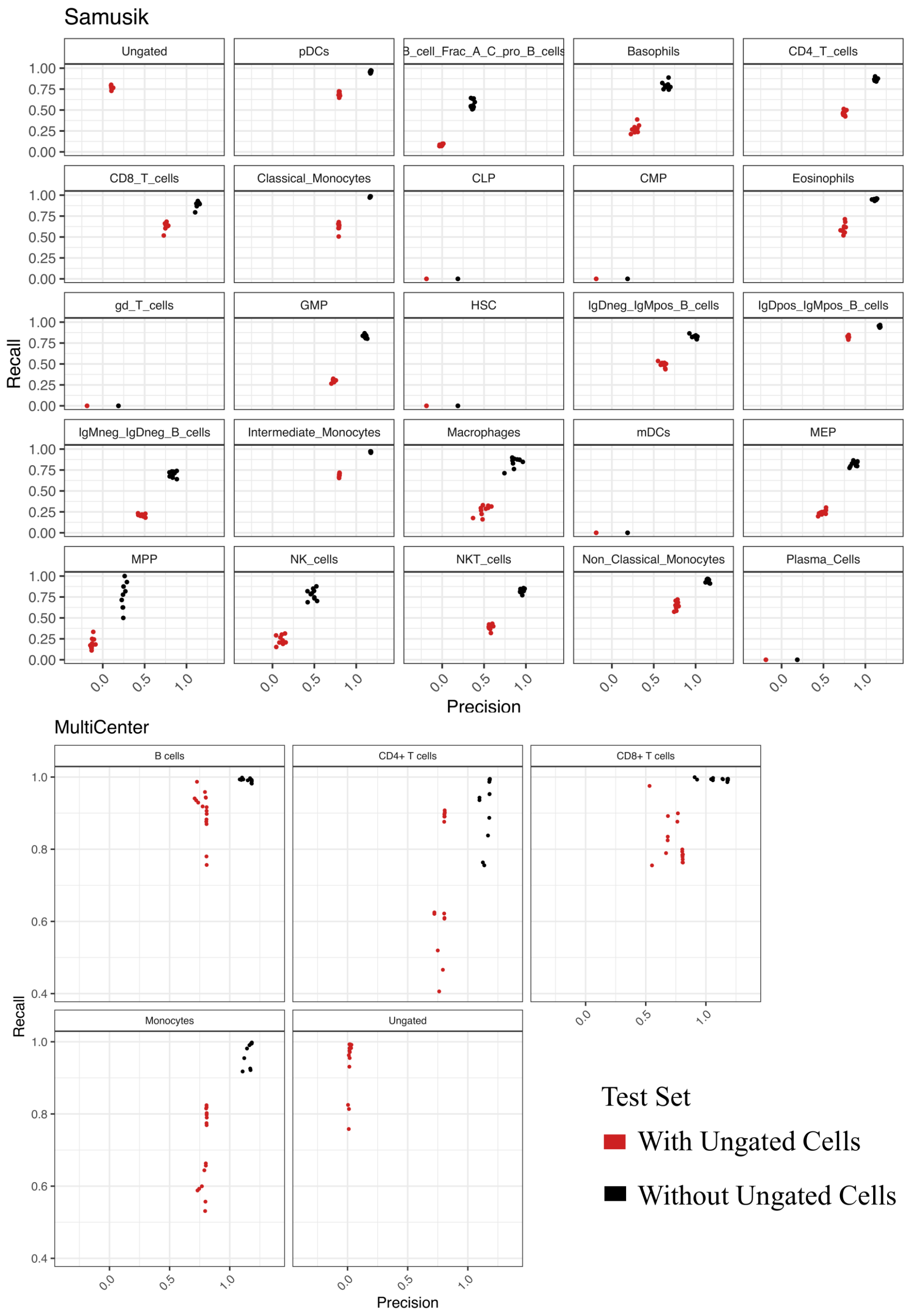


**Figure S3A**. DeepCyTOF. The change in precision and recall score of prediction if ungated cells were excluded from or included in the test dataset. The precision vs recall rate shows that the models' (DeepCyTOF) ability to classify gated cells in each sample decreases when ungated cells are taken into consideration. When ungated cells were included in testing set, for most of the cell types, we observed a low precision rate with a high recall rate, which suggests that most of these ungated cells are misclassified to one of the cell types.

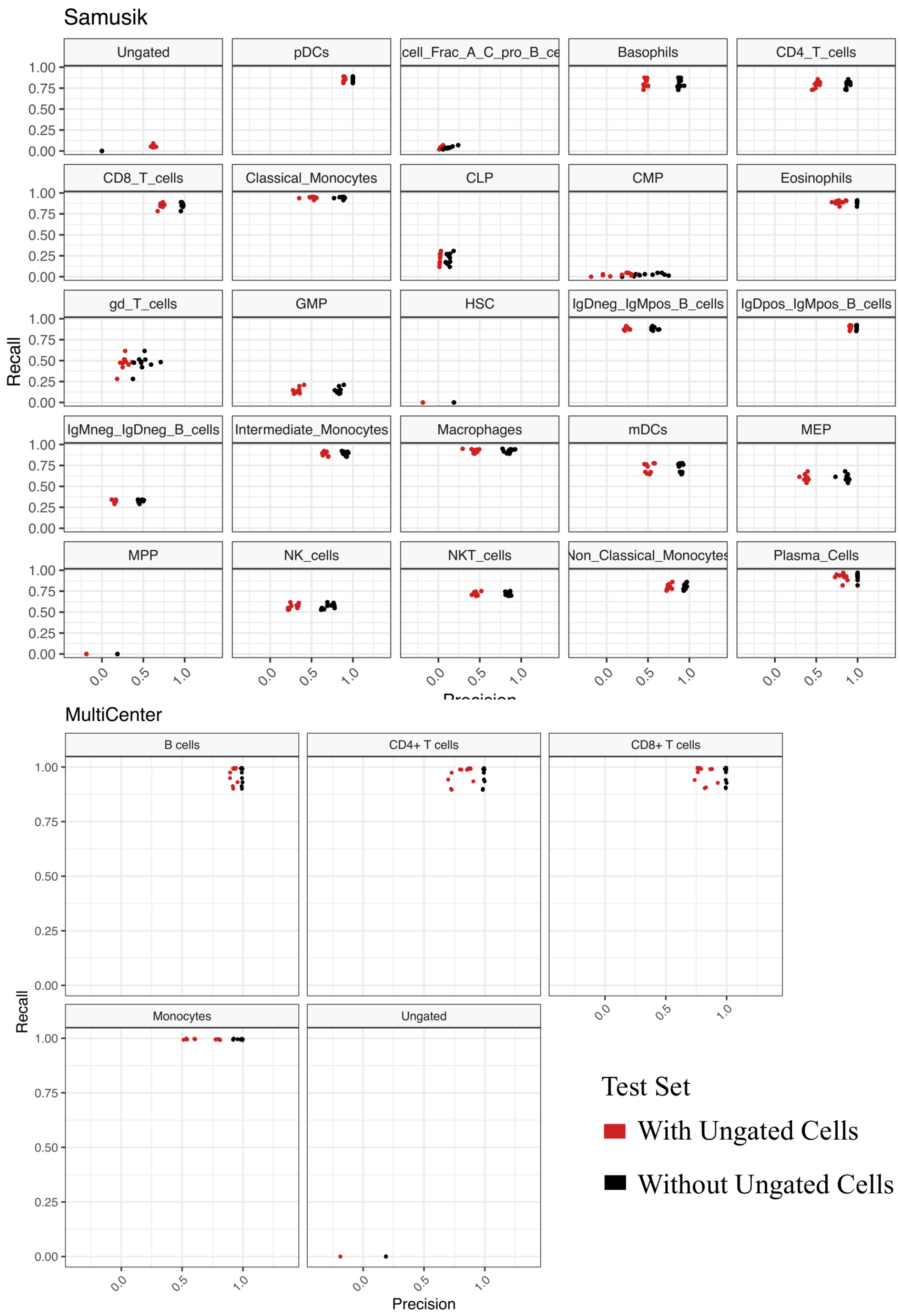


**Figure S3B**. LDA. The change in precision and recall score of prediction if ungated cells were excluded from or included in the testing dataset. The precision vs recall rate shows that the models' (LDA) ability to classify gated cells in each sample decreases when ungated cells are taken into consideration. When ungated cells were included in testing set, for most of the cell types, we observed a low precision rate with a high recall rate, which suggests that most of these ungated cells are misclassified to one of the cell types.

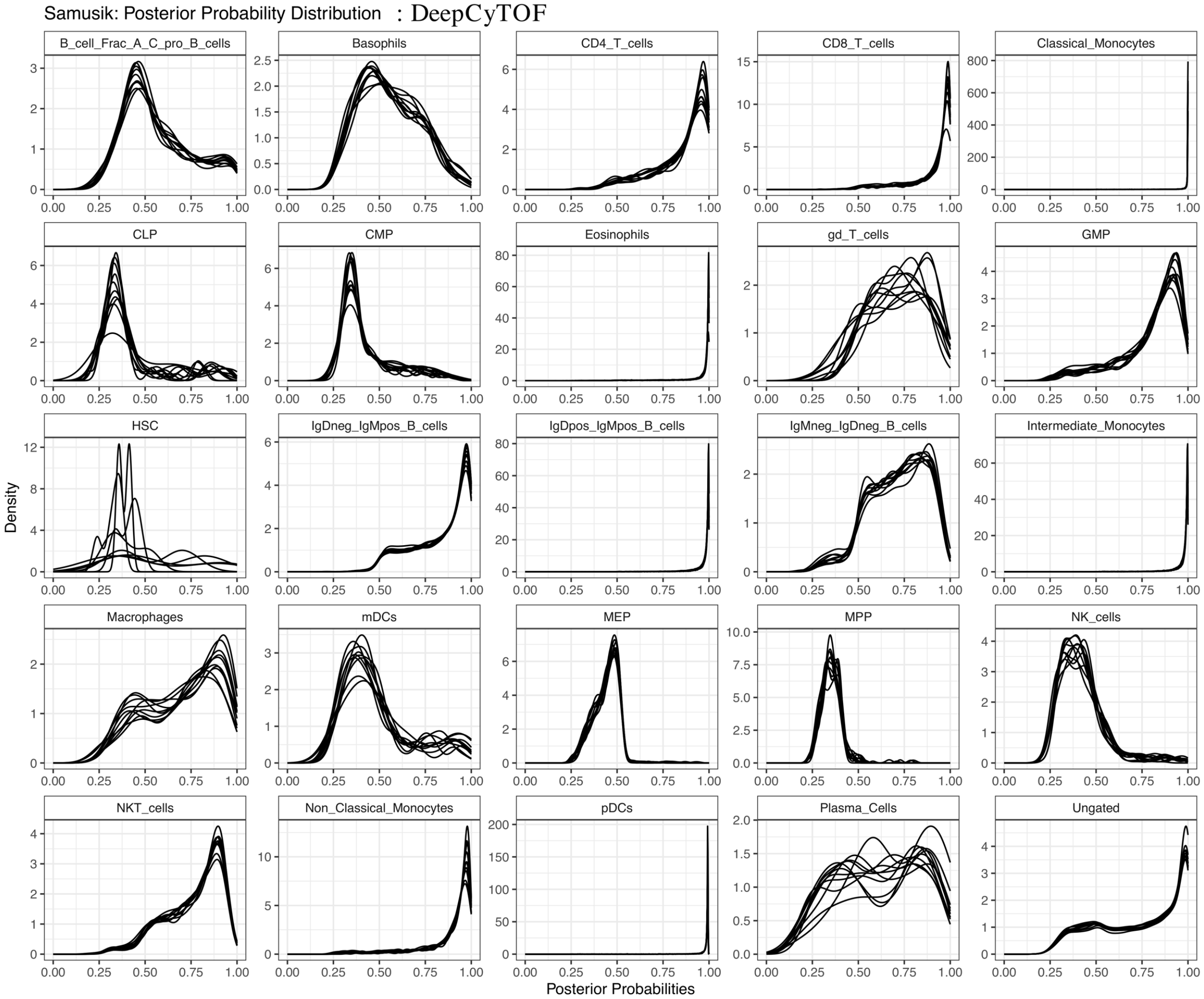


**Figure S4A**. DeepCyTOF on Samusik dataset. For each cell type, the posterior probability distribution observed for cells labeled as the respective cell type after subjecting the dataset to DeepCyTOF.

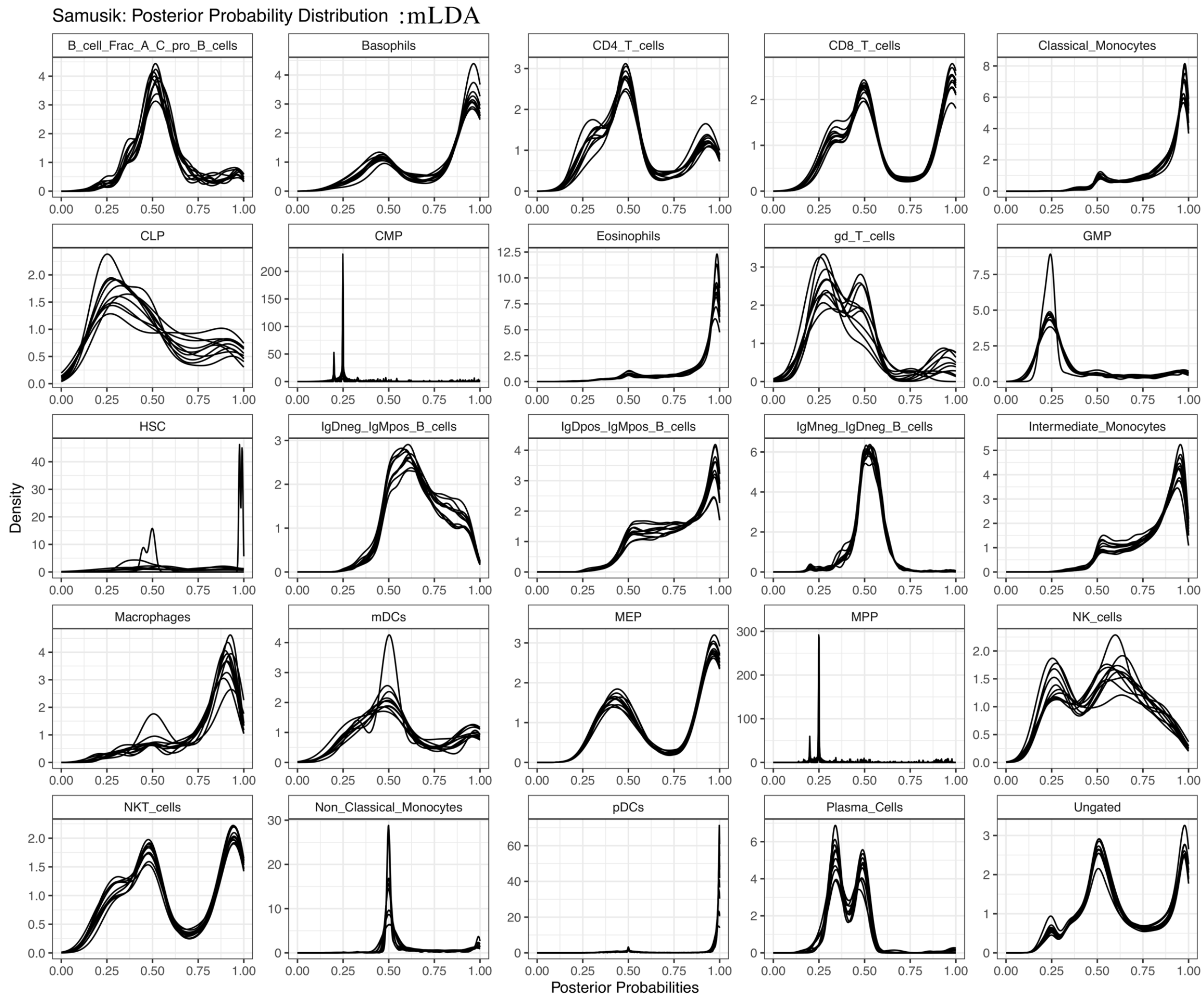


**Figure S4B**. LDA on Samusik dataset. For each cell type, the posterior probability distribution observed for original cell labels after subjecting the dataset to LDA.

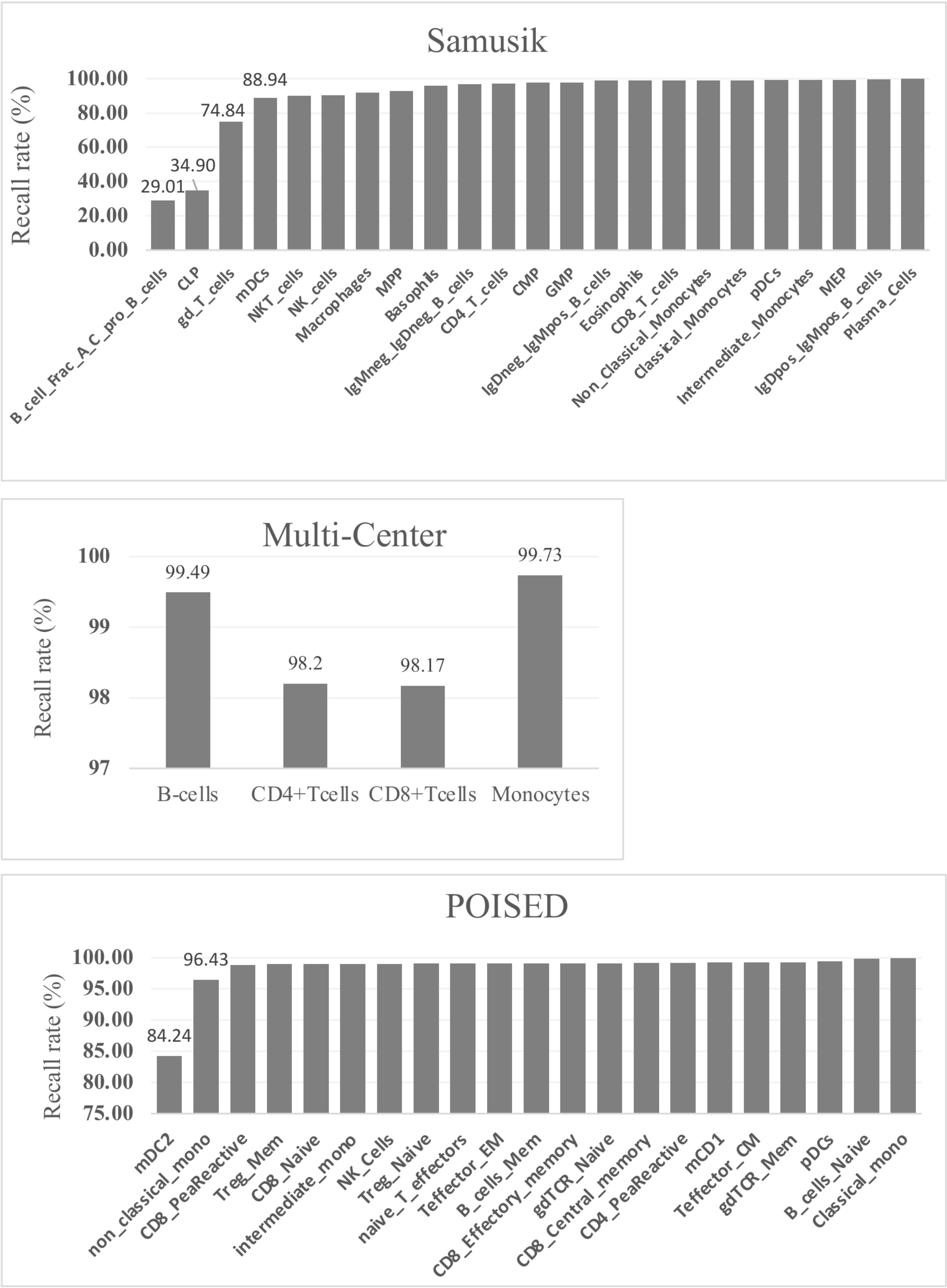


**Figure S5**. Recall rate. The percentage of true positives cells that are the nearest neighbors of their respective set of landmark cells in each cell type, in each dataset.

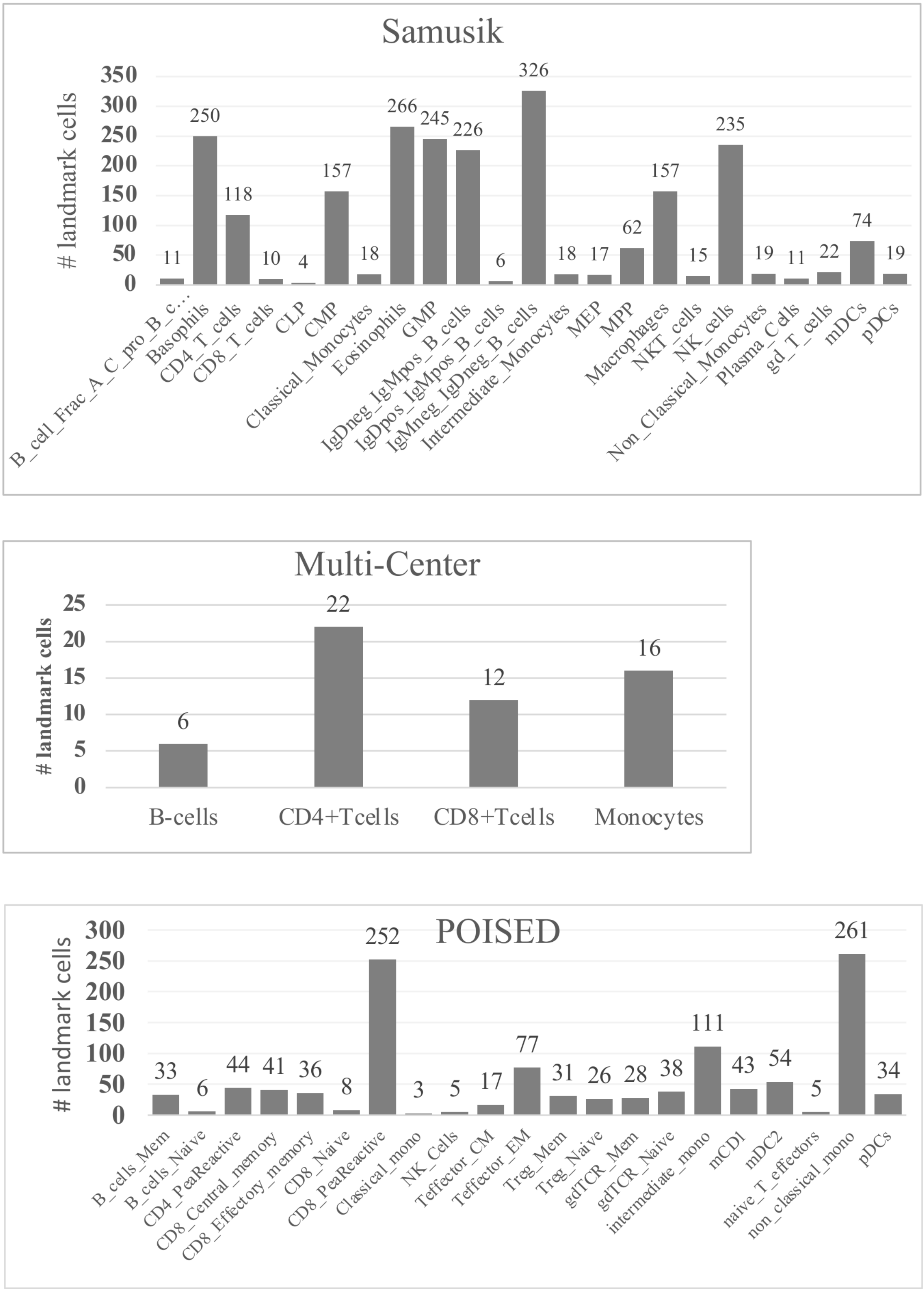


**Figure S6**. Landmark cells. The number of unique landmark cells in each cell type, in each dataset used.

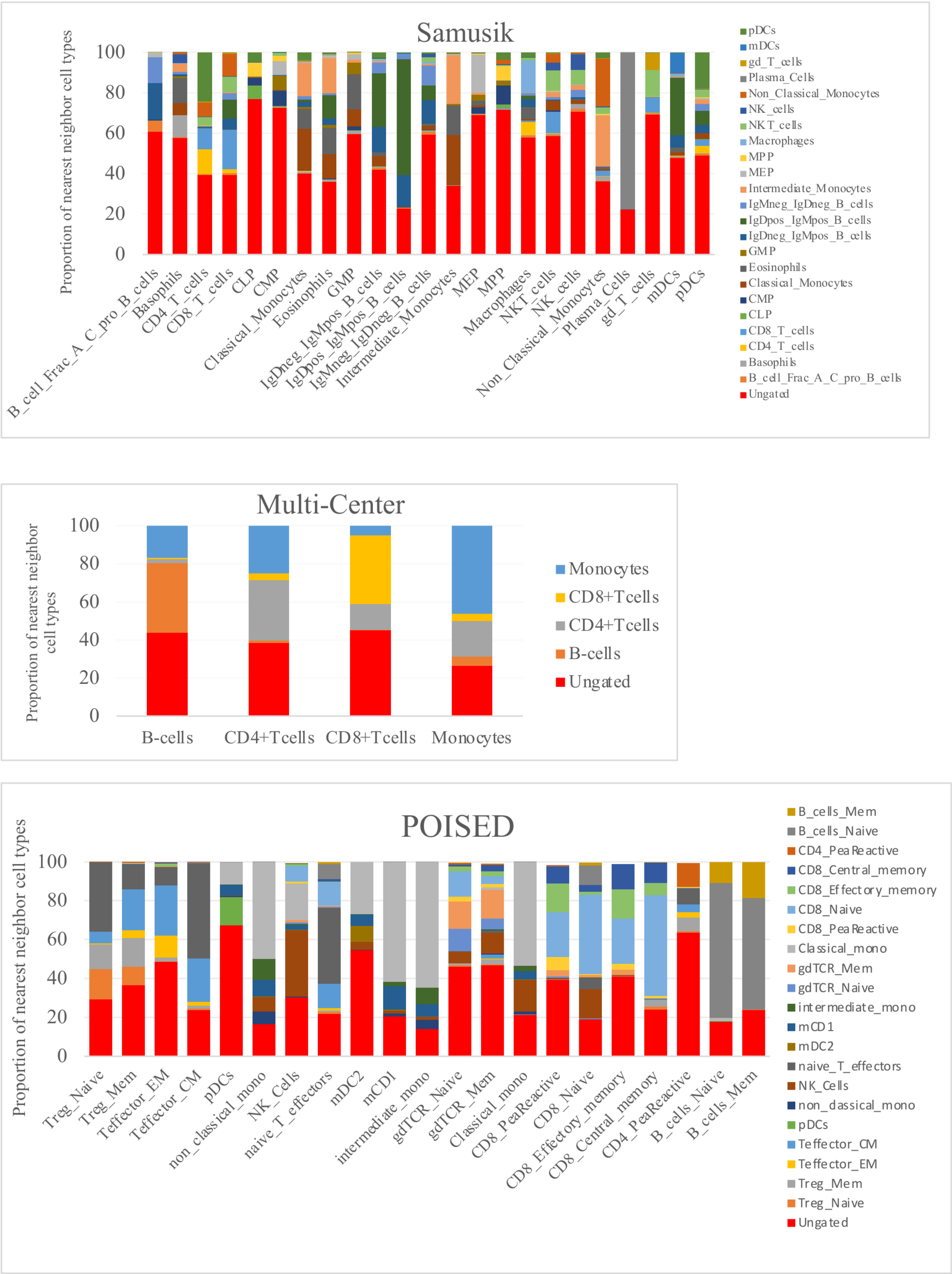


**Figure S7**. Nearest neighbors. The proportion of different cell types in each “cell type specific dataset”. Ungated cells form a significant proportion of the false positive nearest neighbors in most of the cell type specific dataset.

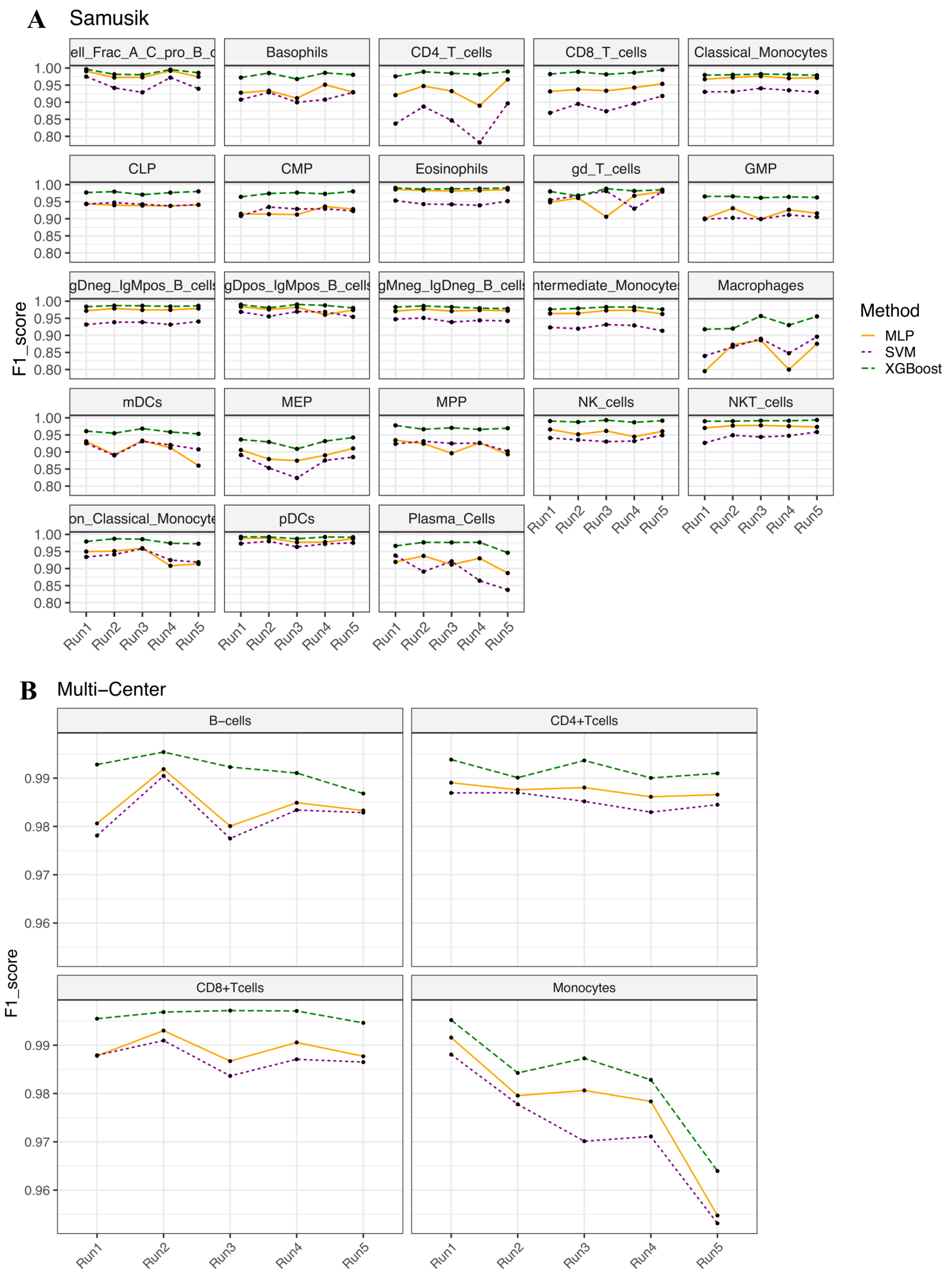


**Figure S8A-B**. F1 accuracy achieved for each cell type with different classification algorithms used in CyAnno in A. Samusik dataset and B. Multi-Class dataset

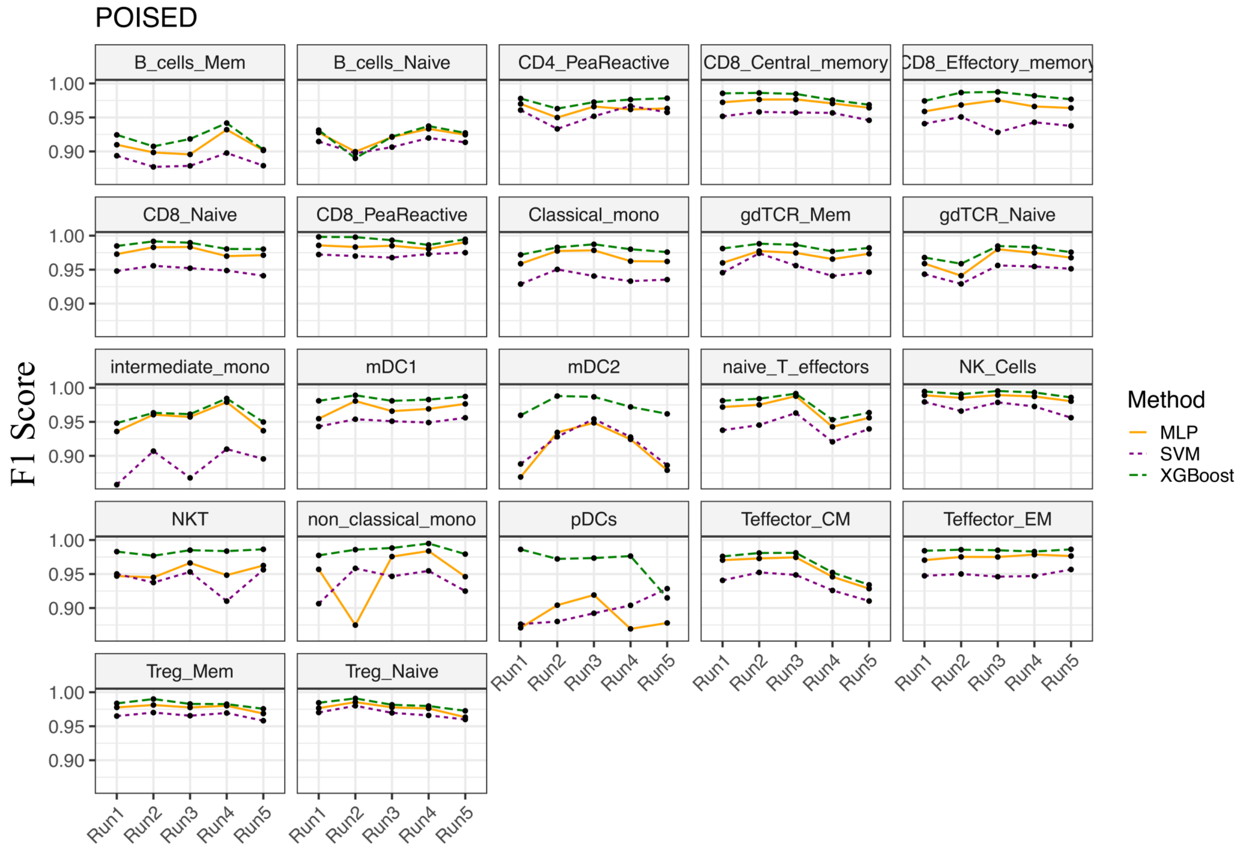


**Figure S8C**. F1 accuracy achieved for each cell type with different algorithms used in CyAnno in POISED dataset

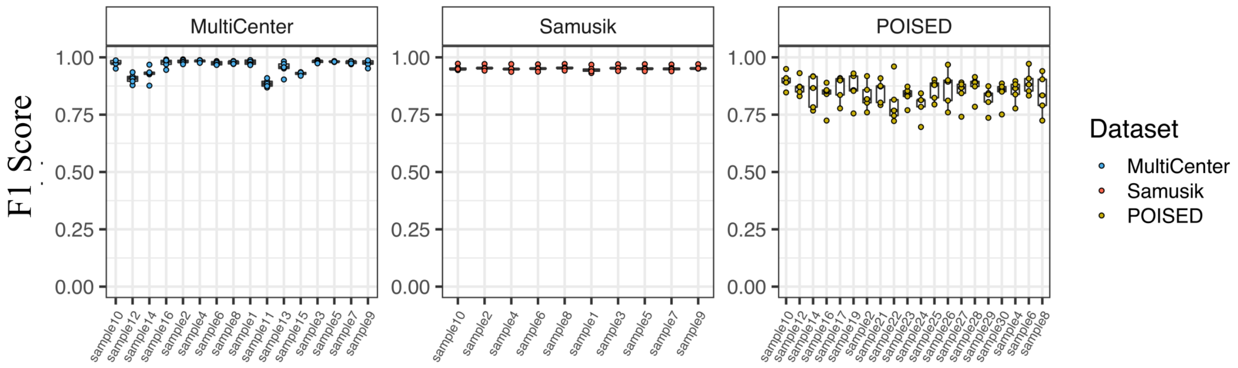


**Figure S9**. F1 Score variability in each sample after 5 different runs with CyAnno. Each run involves varying training samples. For POISED dataset, training and test sets are completely independent, mixed with samples from 7 different batches and 2 different stimulations.

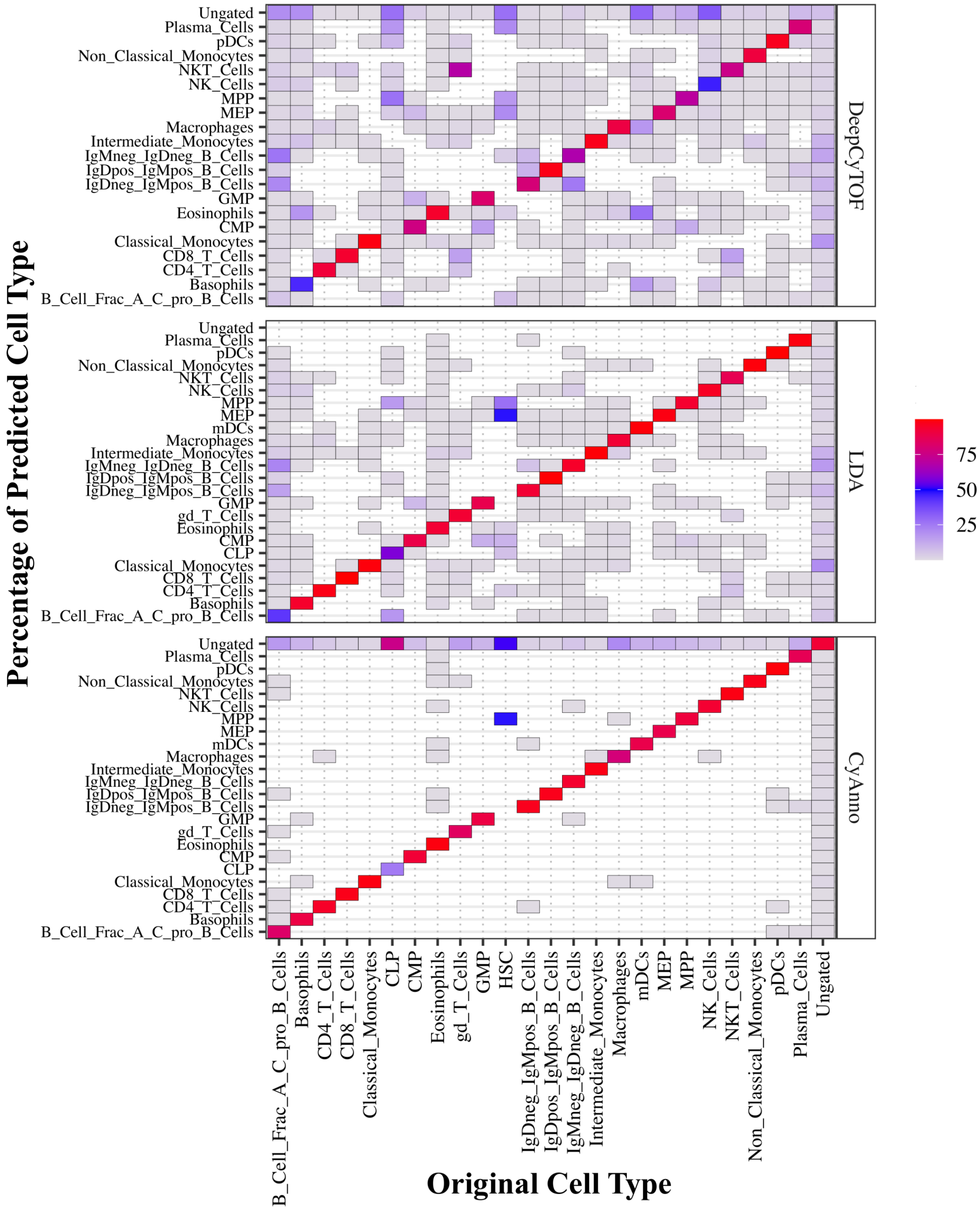


**Figure S10A**. Samusik dataset. For all the cells in each of the original cell types, the composition (as percentage) of different cell type labels predicted by the models is shown. The different cell type labels predicted for a given original cell type suggests that most of the FP prediction in CyAnno corresponds to ungated class of cells. Here, 'Red' tiles represent high percentage of corresponding cell type labels predicted for a given original cell type, whereas blue represent low and white represent that the corresponding cell type label is not predicted for any of the cells from the original cell type.

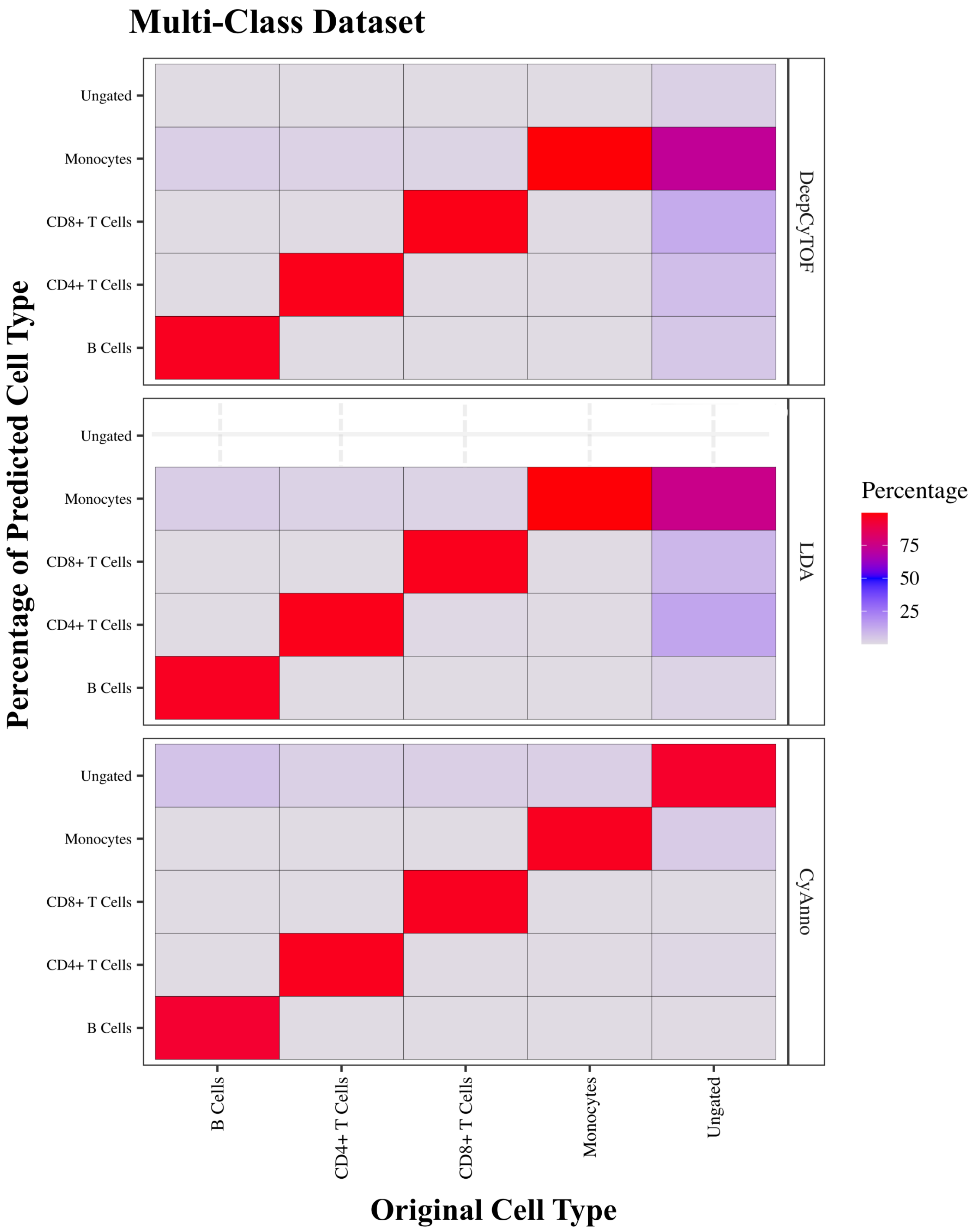


**Figure S10B**. Multi-Class dataset. For all the cells in each of the original cell types, the composition (as percentage) of different cell type labels predicted by models. The composition of different cell type labels predicted for a given cell type reflects large FP prediction. Red boxes represent high percentage of corresponding cell type labels predicted for a given cell type, whereas blue represent low and white represent that the corresponding cell type is not predicted for any of the cells from original cell type.

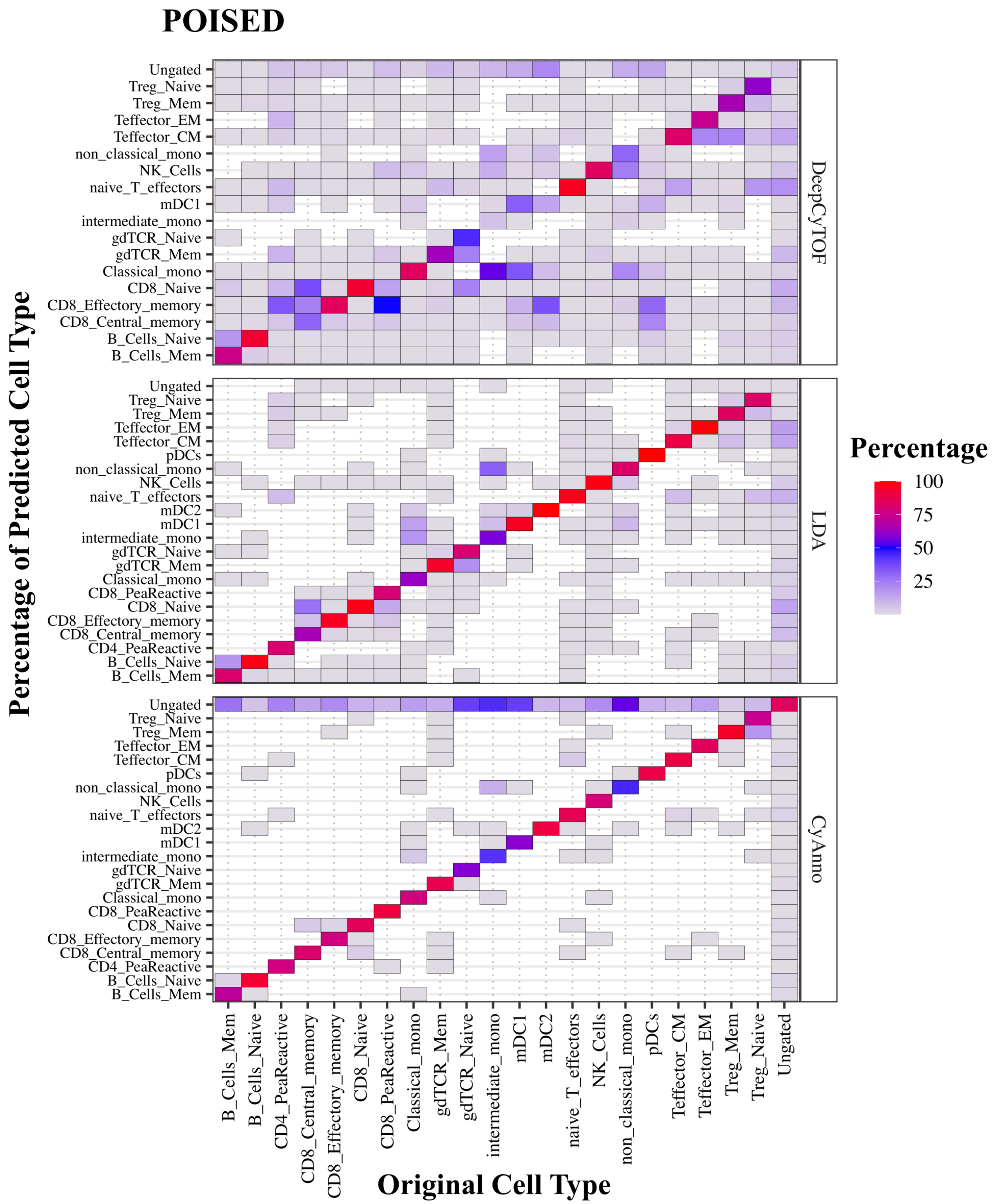


**Figure S10C**. POISED dataset. For all the cells in a given cell type, the composition (as percentage) of different cell type labels predicted by models. The composition of different cell type labels predicted for a given cell type reflects large FP prediction. Red boxes represent high percentage of corresponding cell type labels predicted for a given cell type, whereas blue represent low and white represent that the corresponding cell type is not predicted for any of the cells from original cell type.

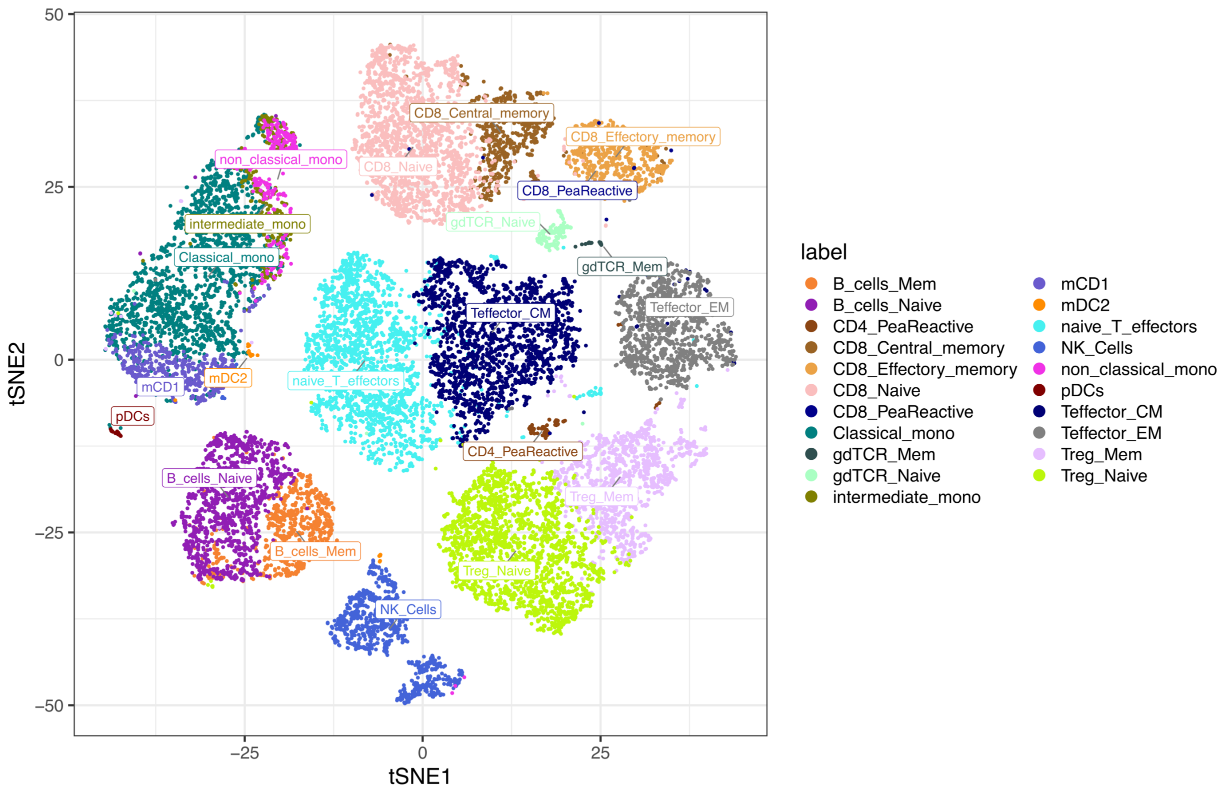


**Figure S11A**. POISED dataset. tSNE visualization for POISED dataset from one sample. Maximum of 2000 cells per cell type was used for plotting the different hand-gated cell types used in this study. For clarity, ungated cells were removed for the dataset to highlight rare/small group of gated cells, e.g. peanut reactive cells (i.e. CD4+ PeaRective and CD8+ PeaReactive). It is observed that even certain gated cell types are difficult to distinguish and share non-linear classification boundaries in a 2D space, e.g. CD8+ Peanut reactive T cells or Memory B cells.

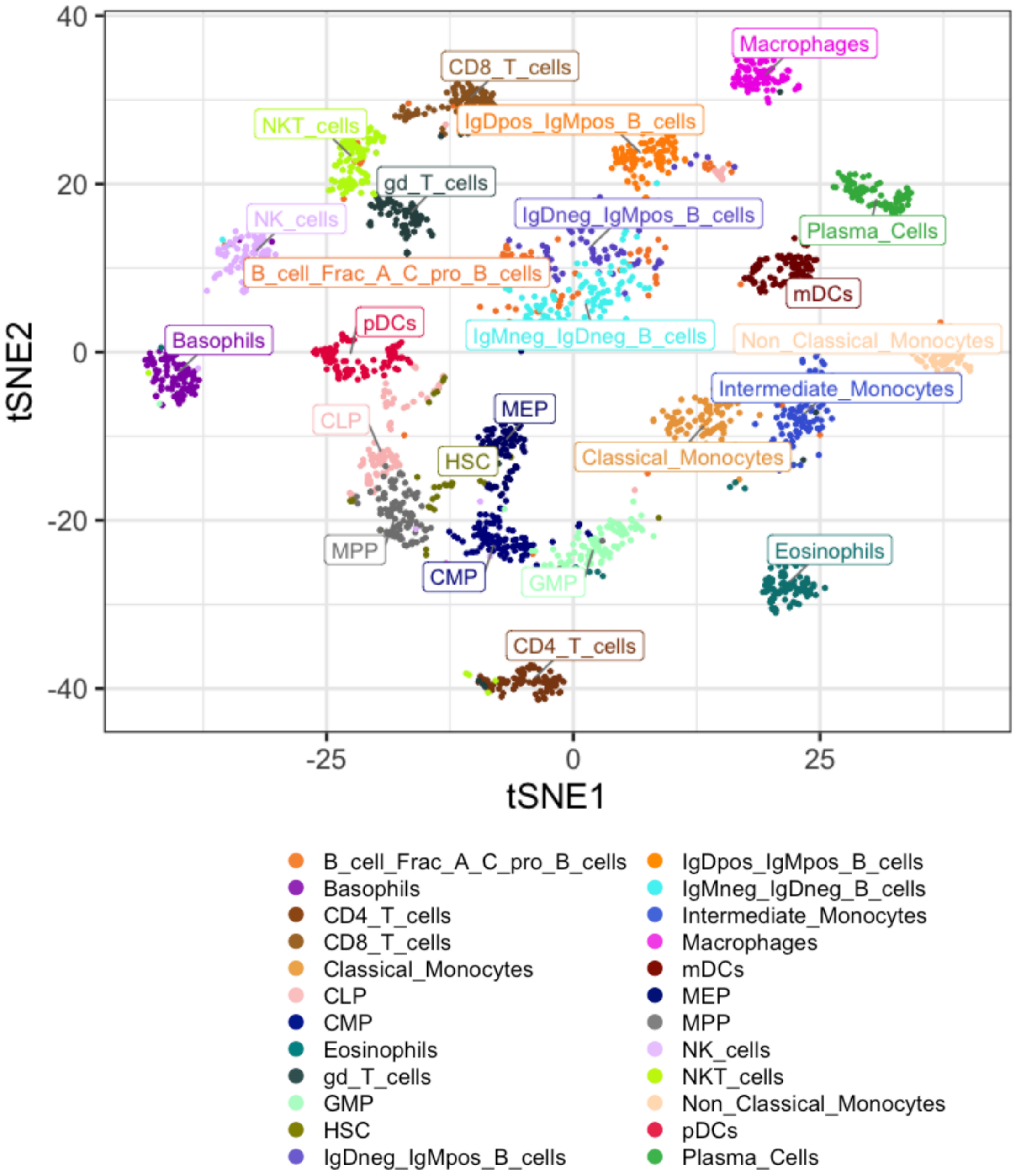


**Figure S11B**. Samusik dataset. tSNE visualization for Samusik dataset. Maximum of 2000 cells per cell type, from all the samples, was used for plotting the different hand-gated cell types used in this study. For clarity, ungated cells were removed for the dataset to highlight rare/small group of gated cells, e.g. HSC.

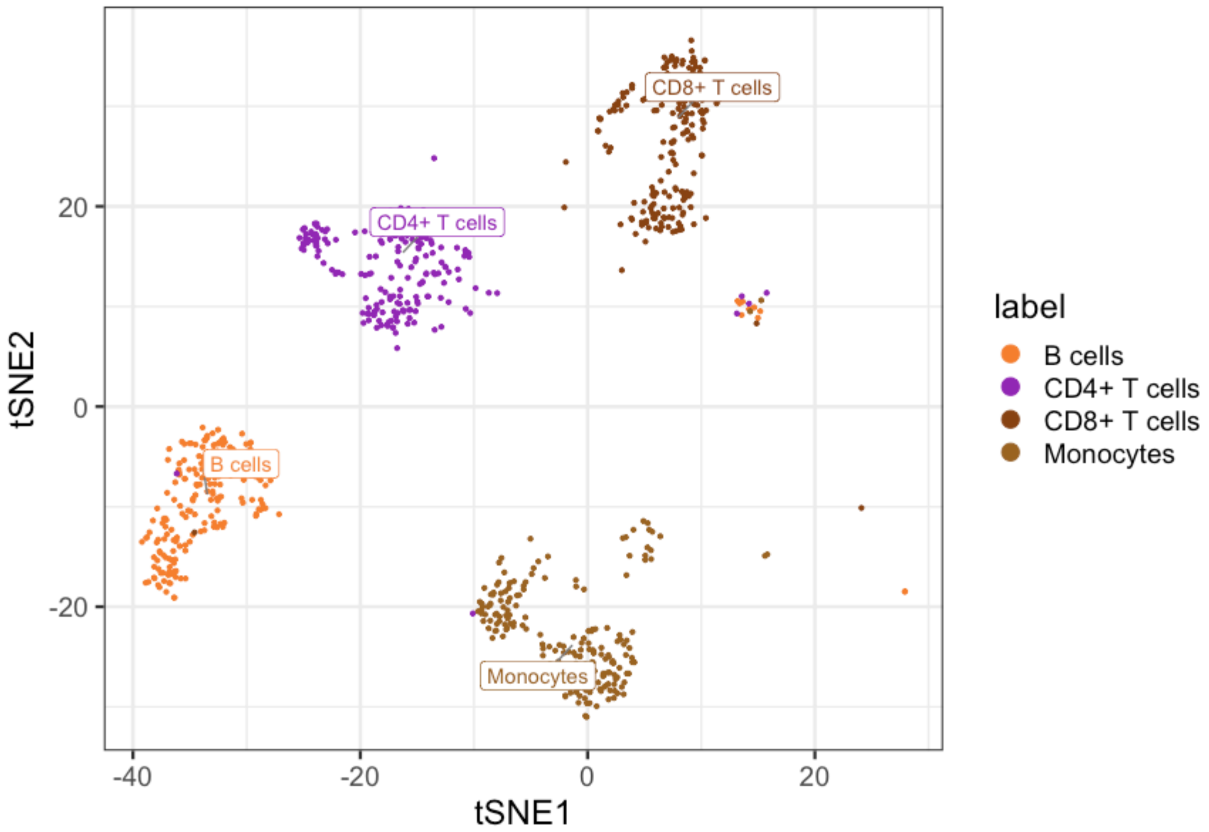


**Figure S11C**. MultiCenter dataset. tSNE visualization for MultiCenter dataset. Maximum of 2000 cells per cell type, from all the samples, was used for plotting the different hand-gated cell types used in this study. For clarity, ungated cells were removed for the dataset.

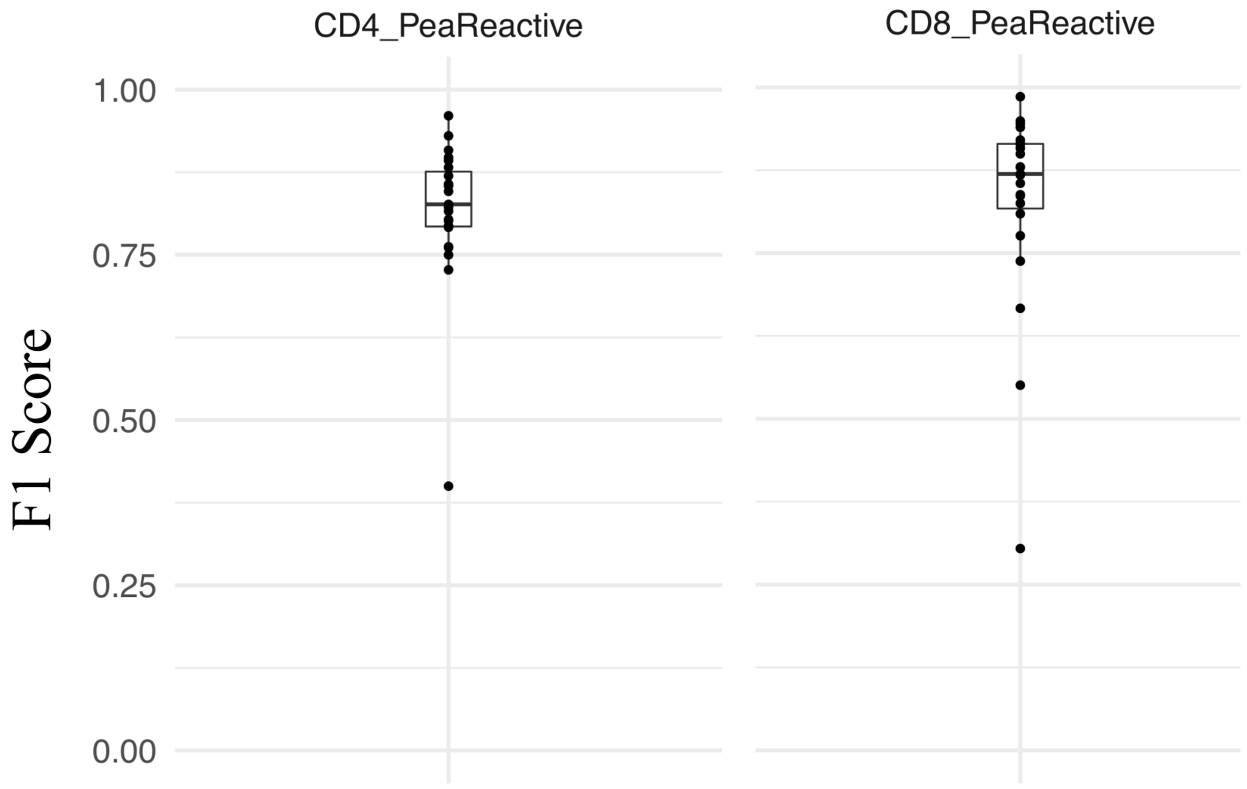


**Figure S12**. The F1 score observed with CyAnno, when the algorithm was trained to predict only the CD4 peanut reactive cell type or only the CD8 peanut reactive cell type, respectively in a different run. The predicted F1 score reflects the accuracy by which CD4 peanut reactive cells and CD8 peanut reactive cells were predicted in the independent test dataset with 20 samples.

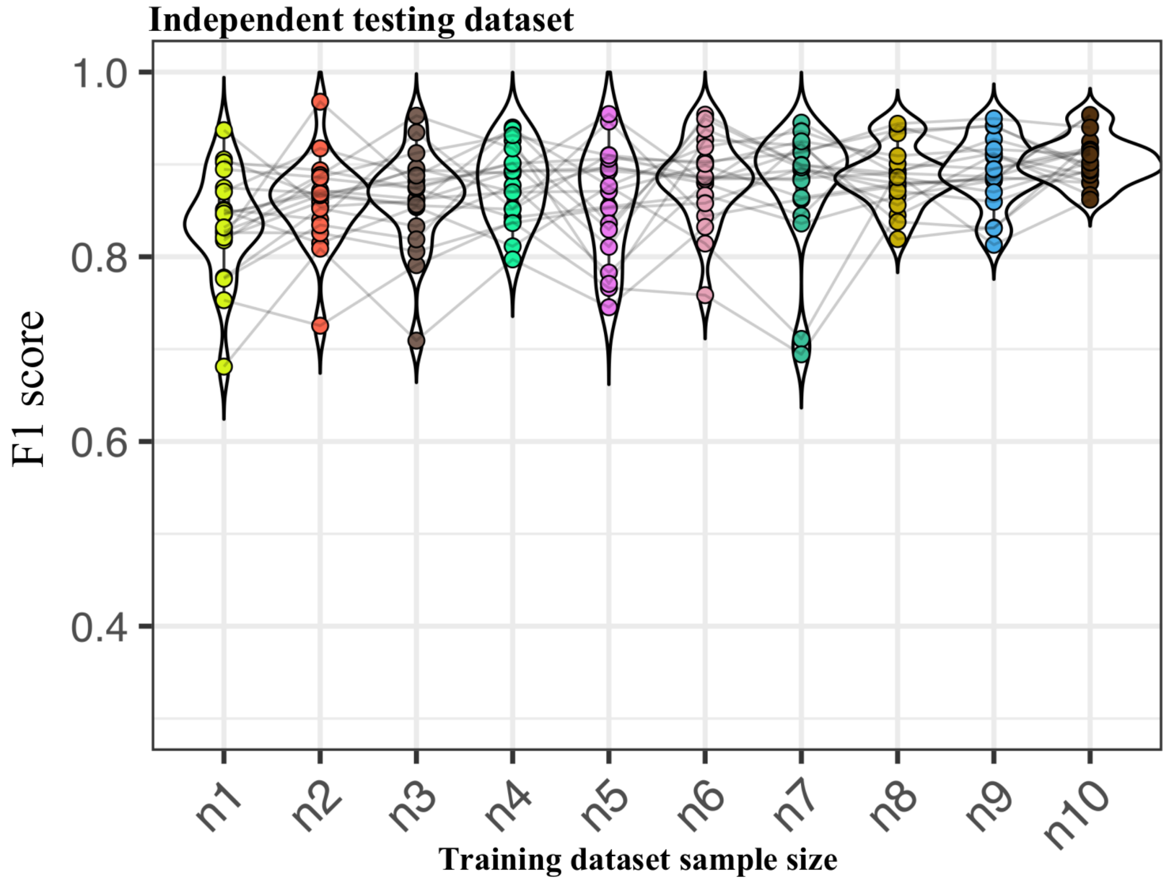


**Figure S13**. CyAnno prediction with sample size variation. The F1 score per sample in independent testing set when CyAnno was executed with varying number of samples (*n*) in training set, ranging from *n*=1 to *n*=10.

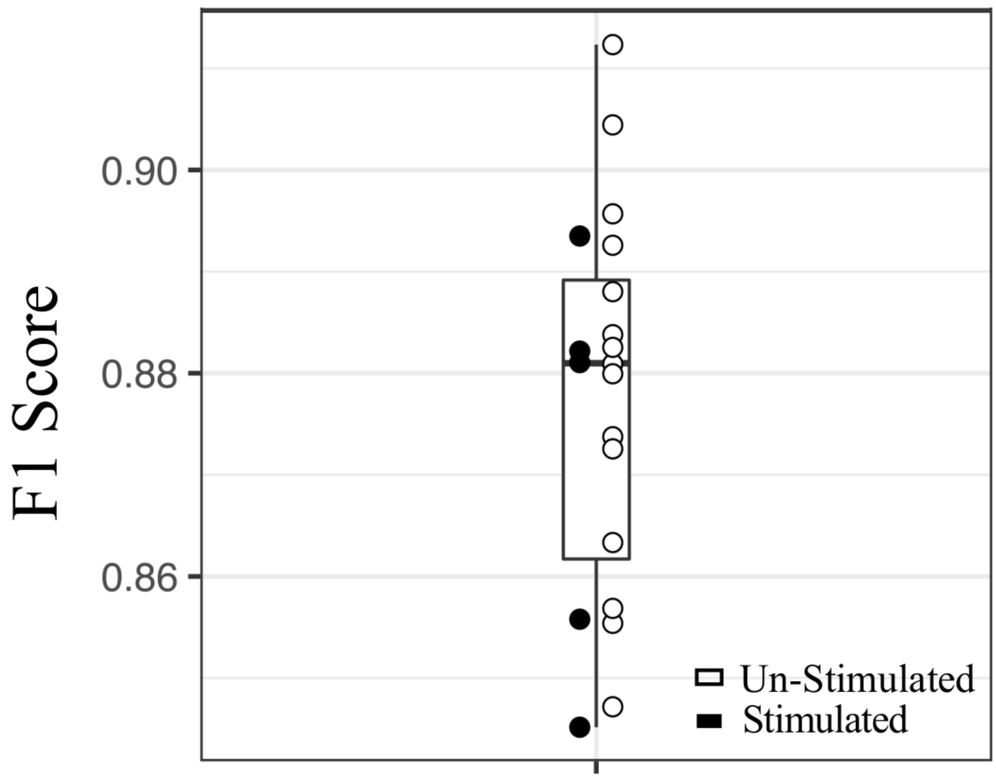


**Figure S14**. The F1 score observed with CyAnno in test set samples (n=20) colored by their stimulation status with training set composed of only peanut stimulated samples.

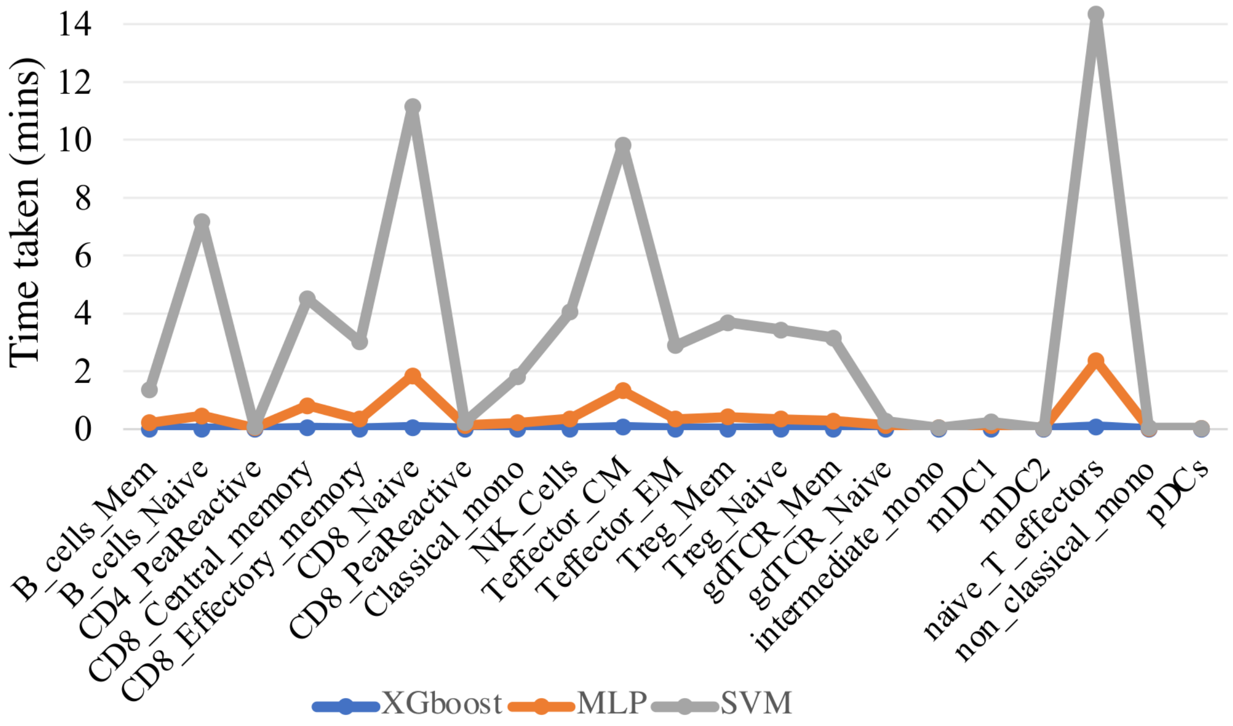


**Figure S15**. Time taken by different ML classifiers used in CyAnno to train and build the hyper-parameter optimized CTSM.
